# Generalizable Finger Movement Decoding from Intracranial Recordings Across Static and Dynamic Actions

**DOI:** 10.64898/2026.03.27.714722

**Authors:** Eva Calvo Merino, Qiang Sun, Yihan Wu, Jiawei Liao, Yuxin Quan, Tao Chang, Mwibwe Mulenga, Yanhui Liu, Qing Mao, Yuan Yang, Jiayuan He, Marc M. Van Hulle

**Author notes:** These authors contributed equally to this work (Eva Calvo Merino, Qiang Sun, Yihan Wu). Shared corresponding authors (Eva Calvo Merino, Yuan Yang, Jiayuan He).

## Abstract

Reliable decoding of finger movements requires brain-computer interfaces (BCIs) that generalize across the diverse static and dynamic actions performed in daily life. Static postures and dynamic movement sequences rely on partially distinct neural representations, and the balance between these components varies widely across tasks. Yet most BCI decoding pipelines are optimized for a narrow task domain, limiting their ability to transfer to new scenarios. Here we show that standard intracranial decoding approaches show notable performance drops when applied to previously unseen finger tasks with different static-dynamic compositions, and that generalization critically depends on multiple elements of the decoding pipeline. High gamma features and short temporal windows yield the most robust cross task transfer. Although nonlinear models perform best when examples of all tasks are represented during training, linear decoders generalize better to novel, particularly dynamic, actions. We identify a consistent static-dynamic structure across tasks that may provide a useful basis for improving generalization. Finally, we show that anatomical heterogeneity between tasks also constrains generalization. Together, these results establish design principles for BCIs that generalize reliably across the rich repertoire of human finger movements.

## Introduction

Dexterous finger movements are essential to most daily activities, providing humans with great flexibility in interacting with the environment and enabling the creation and manipulation of tools ^1^. This central role makes fingers a primary target for brain-computer interfaces (BCIs) ^2^. Finger BCIs hold significant promise: in the near term, they could restore dexterity in individuals with paralysis^3^, while in the longer term, they may enhance motor capabilities for the general population ^4^. Most finger BCIs rely on intracranial recordings, as they provide the signal quality and spatial resolution needed for finger decoding^5–8^, which are difficult to achieve with non-invasive recordings ^9,10^. In particular, many studies use electrocorticography (ECoG), where electrodes are placed on the cortical surface to capture stable neural activity with high signal quality .

Over the past decade, ECoG studies have shown promising decoding performance of individual finger movements^11–17^. Despite these advances, current finger BCIs remain limited, particularly in their ability to generalize across the diverse range of human finger actions. This lack of adaptability presents a major challenge for restoring versatile hand function. From a motor-control perspective, finger actions can be divided into two main groups: (i) static actions (isometric), where muscles generate force without changing length to maintain a fixed posture (e.g., pointing or holding an object), and (ii) dynamic actions, where finger position changes continuously, involving changes in muscle length (e.g., typing) ^18–20^. Ideally, BCIs should work smoothly across both types of actions. However, prior work indicates that these actions differ in their neural representations. Studies using ECoG^21–24^, microelectrodes^24,25^, and non-invasive recordings^26–29^ consistently report stronger cortical activation during dynamic movements compared to static postures, across multiple frequency bands ^22,23,28,29^ and in both motor ^20,21^ and sensory ^24–26^ areas. Moreover, these two actions have been associated with partially distinct anatomical regions^21,25^, with more spatially localized activation during static postures and broader activation during dynamic movements. These observations suggest that a BCI trained on one type of action may not generalize well to the other ^23^.

Nevertheless, most ECoG-based finger BCI research has focused on a single task ^11–15,17^: continuous, repetitive flexion-extension of individual fingers, largely because this is the task considered in publicly available datasets ^30,31^. These datasets have driven optimization of BCI decoding pipelines, shaping both feature design and decoder selection. From a feature perspective, low-frequency phases and high-gamma power are widely considered the most informative ^5,14^, typically extracted from a one-second lookback window preceding the time sample being decoded ^32–34^. From a modeling perspective, both linear and non-linear decoders have been explored, with neural networks achieving the highest performance ^11–13,17,34,35^. However, these findings are based on highly dynamic tasks. Will the knowledge transfer to tasks with different static-dynamic action compositions?

In this study, we investigate decoding strategies for BCIs that generalize across static and dynamic finger actions using high-density ECoG recordings from four subjects, with electrodes placed over primary motor and somatosensory cortices. We analyze two tasks: one dominated by static postures (sustained task; Fig. 1a,d up) and another dominated by dynamic movements (continuous task; Fig. 1b,d down). We evaluate performance in two scenarios: a seen case, where decoders are trained and tested on the same task, and an unseen case, where decoders trained on one task are transferred to the other. We focus on the unseen scenario because future finger BCIs must be able to predict movements not exposed during decoder training, thereby simulating the emergence of new actions during natural interactions.

**Fig. 1.**
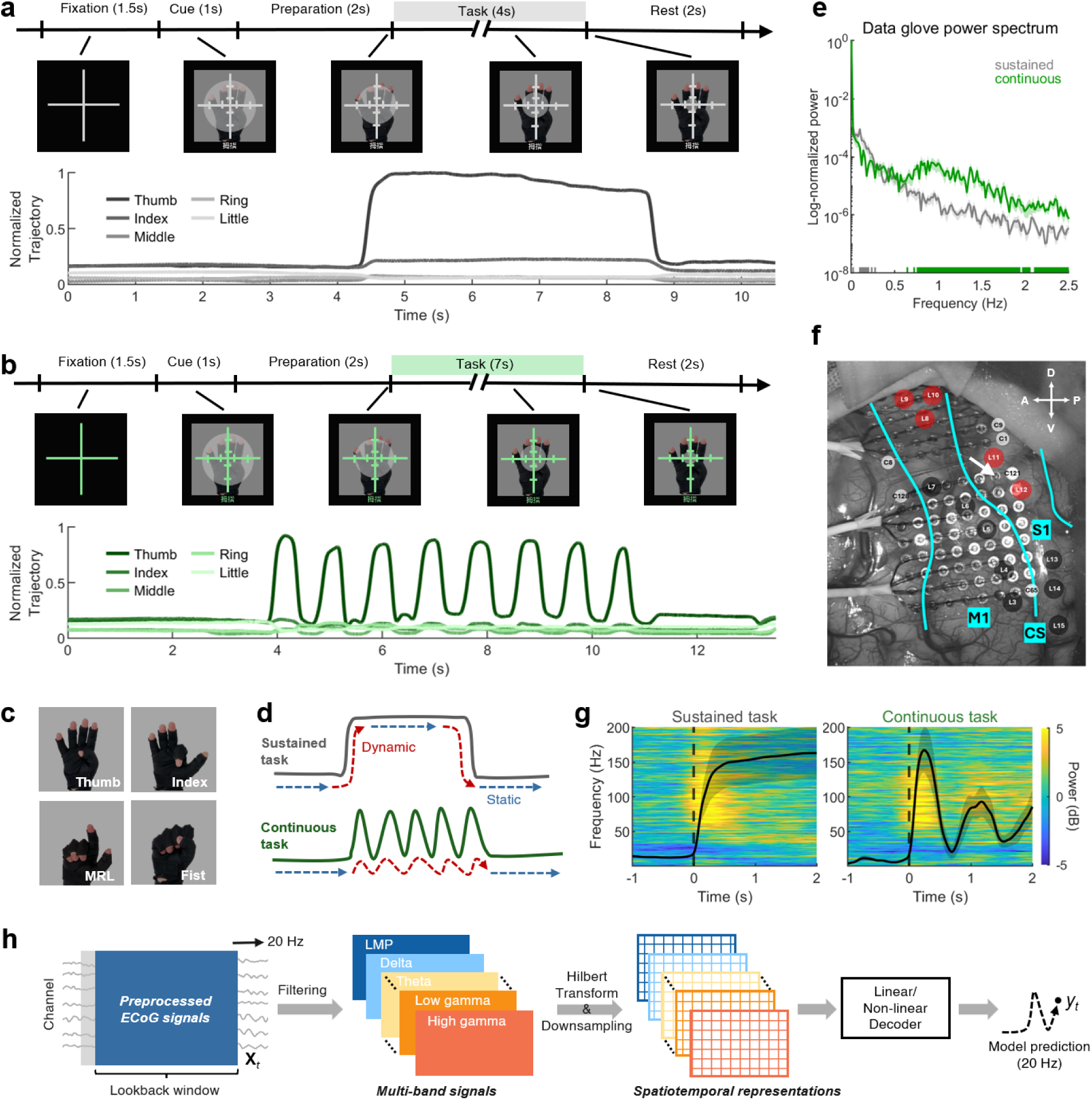
Graphical overview of the methods. a, **b.** Example thumb-movement trial of the sustained (a) and continuous (b) finger-movement tasks. Top: trial timeline with visual cues presented to participants. Bottom: normalized finger trajectories recorded with a data glove (one sensor per finger). The longer task period in the continuous condition allows multiple flexion-extension cycles. **c.** The same four finger movements were performed in both tasks (MRL: middle-ring-little). **d.** Illustration of static and dynamic actions for example trajectories in each task. **e.** Average power spectrum of data glove trajectories across all subjects and fingers for both tasks (sustained: grey; continuous: green). Frequencies with significant differences between tasks (Wilcoxon paired test, FDR-corrected) are indicated. **f.** ECoG grids (8×8; 128 channels) placed over sensorimotor cortex of subject 2. Sites where electrical stimulation evoked finger movement or sensation are marked in red; other responsive sites are shown in grey (see Supplementary Fig. S1 for a detailed representation of all subjects). Cyan lines demarcate the central sulcus (CS), primary motor cortex (M1), and primary sensory cortex (S1). Orientation labels denote anterior (A), posterior (P), dorsal (D), and ventral (V). The arrow marks the example channel shown in **g**. **g.** Mean data glove trajectory across all movement trials (± variance) superimposed on the spectrogram and aligned to movement onset, for an example sensory channel (arrow in **d**). Spectrograms are baselined to activity in the interval -2 to -1.5 s. **h.** Schematic of preprocessing and decoding.

Our analysis examines four key components of the BCI decoding pipeline: (i) neural features, (ii) decoder architecture, (iii) action types (static versus dynamic), and (iv) anatomical recording site. We find that each of these components has a substantial impact on generalization. Some neural features, such as low frequency phases or beta- and low gamma-band activity, perform well in the seen case but generalize poorly, whereas high-gamma activity is robust across both scenarios. Decoder design also plays an important role. Models that rely on short temporal windows (< 250 ms) generalize better than the one-second windows commonly used in previous studies, likely because they emphasize movement-related activity rather than task structure. The choice of architecture further affects performance, while nonlinear models outperform linear models in the seen case, this advantage disappears in the unseen condition, where linear models slightly outperform overall. Interestingly, this effect action-dependent: linear models generalize better for dynamic movements, whereas nonlinear models remain superior for static postures. Finally, anatomical differences between the tasks also reduce cross-task generalization. Together, these findings highlight critical design choices for BCI decoding pipelines that generalize across static and dynamic actions.

## Results

Participants performed two finger-movement tasks, sustained and continuous, while finger trajectories were recorded with a data glove (Fig. 1a,b). Both tasks included the same four finger movements (Fig. 1c) and were completed within the same session, with trials cued in random order, ensuring that comparisons are not affected by session timing. The two tasks differ in their static-dynamic composition (Fig. 1d) and in the frequency content of the movement (Fig. 1e, based on data glove trajectories): the sustained task is dominated by movements below 0.5 Hz, whereas the continuous task also contains movements above 0.5 Hz due to its repetitive flexion-extension cycles. Simultaneously, we recorded high-density ECoG activity using two 8×8 grids placed over sensorimotor cortex (Fig. 1f). Example spectral activity from one sensory channel shows the typical movement-related decrease in low-frequency power and increase in high-gamma power ^36^ (Fig. 1g). In the sustained task, high-gamma activity peaks near movement onset, whereas in the continuous task activity persists throughout dynamic flexion-extension cycles.

We trained two decoding models, Partial Least Squares (PLS) and Long Short-Term Memory (LSTM) network, which we chose as representative linear and nonlinear approaches widely used in neural decoding ^11,17,37^, to regress the finger trajectories from the ECoG signals in a causal manner (Fig. 1h). The signals were originally sampled at 2,000 Hz and were downsampled to 20 Hz after feature extraction to match the sampling rate of the dataglove recordings.

### High-gamma features provide the strongest generalization

We first assess how different neural features support generalization to the unseen task (Fig. 2). We compare band-limited power features derived from the Hilbert envelope, the low-frequency amplitude (LMP, Local Motor Potential) derived from low-pass filtering, and a combined high-gamma + LMP feature set, which has been reported as the optimal feature combination in previous ECoG finger-movement decoding studies^5,14^.

**Fig. 2.**
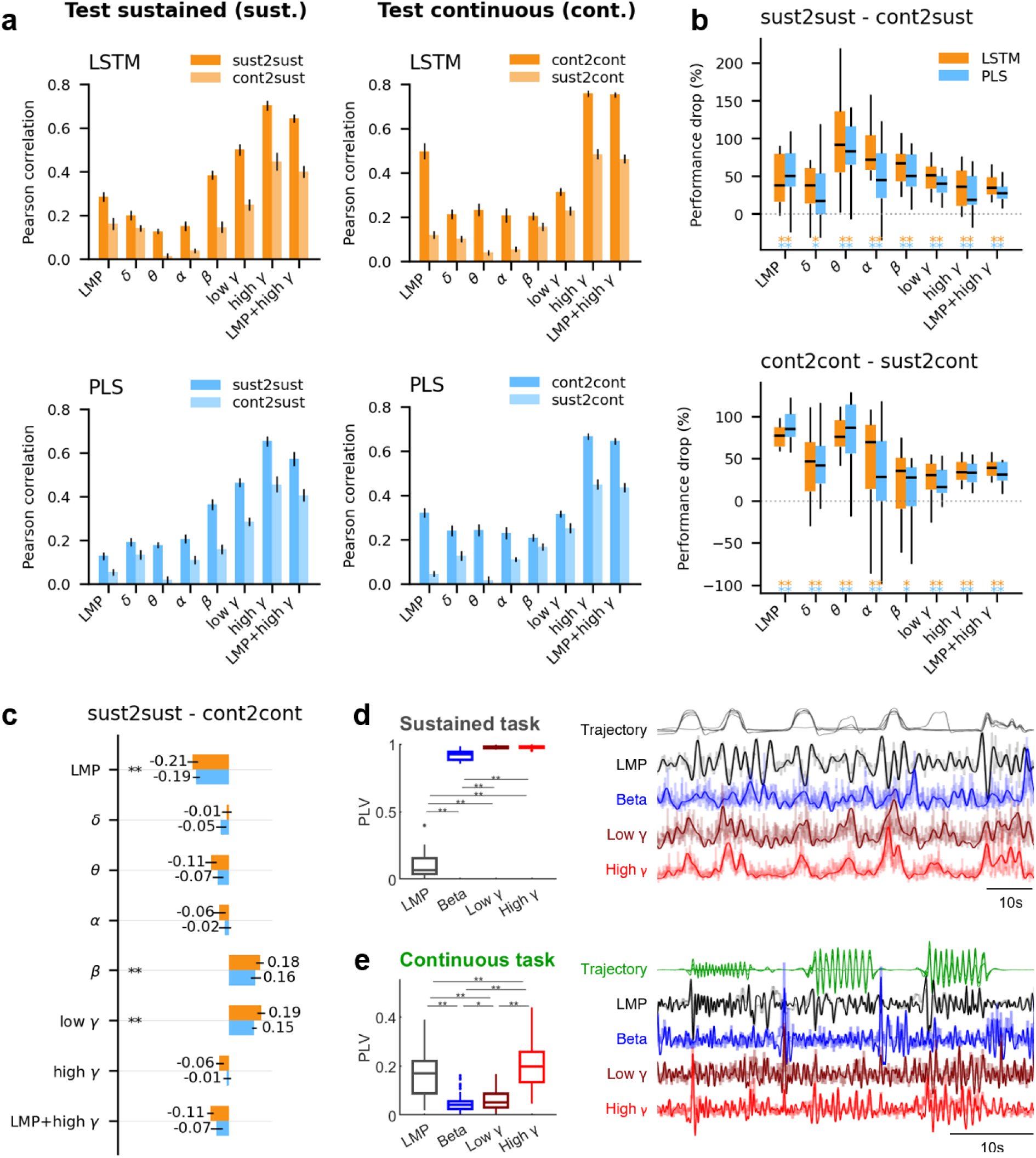
Frequency band contributions to decoding. **a.** Pearson correlation between dataglove trajectories and the predictions of each model (orange: LSTM; blue: PLS) for all features in the seen and unseen conditions. Bars show the mean (± SEM) across all subjects and fingers (n=20). High-gamma significantly outperforms all other features (Wilcoxon paired test, FDR-corrected, p < 0.005), except when compared with high-gamma + LMP in the continuous test condition (p > 0.05). **b.** Performance drop in the unseen condition relative to the seen condition, based on the correlations shown in panel a. All features exhibit a significant drop in performance for both models. **c.** Difference in correlation between the sustained and continuous tasks (seen condition only), illustrating task-specific feature performance. Bars show the mean (± SEM) across all subjects and fingers. **d.** PLV between neural features and dataglove trajectories for the sustained task (< 0.5 Hz; frequency range defined based on Fig. 1e), computed using the top five channels contributing most to the PLS predictions. Each box represents 100 values (5 channels x 4 subjects x 5 fingers). An example of the filtered trajectories and corresponding feature traces is shown on the right (subject 4). **e.** As in panel d, but for the continuous task (0.5–3 Hz; frequency range defined based on Fig. 1e). In panels b, d, and e, boxes show the median and the interquartile range, whiskers show the most extreme non-outlier values (within 1.5×IQR), and outliers are plotted as dots in panels d and e (not displayed in panel b for clarity). Statistical significance is assessed with the Wilcoxon signed-rank test (*p < 0.05, **p < 0.005; FDR corrected).

Across both decoding models, high-gamma yields the highest decoding performance in both the seen (sust2sust, cont2cont) and unseen (sust2cont, cont2sust) conditions (Fig. 2a). High-gamma significantly outperforms all other features (*p <* 0.005, FDR-corrected), except when compared with high-gamma + LMP in the continuous test condition. This aligns with the common understanding that high-gamma activity is the most informative feature for movement-related ECoG decoding ^38^.

All features exhibit a performance drop when evaluated on the unseen task relative to the seen task (Fig. 2b). Low and mid frequency bands (below low-gamma) show the largest drop, a pattern consistent across both decoders. Interestingly, some features show strong task-specific performance, which limits their ability to generalize across tasks (Fig. 2c). LMP performs significantly better for the continuous task than for the sustained task, whereas beta and low-gamma perform better for the sustained task. Although these features are often considered useful in intracranial motor BCIs^38^, our results indicate that they are not well suited for cross-task generalization.

To understand this task-specific behavior, we examine the coupling between each neural feature and the finger-movement trajectories using the phase-locking value ^39^ (PLV; Fig. 2d,e). For each task, neural features and dataglove trajectories are filtered within the task-relevant frequencies (<0.5 Hz for sustained; 0.5–3 Hz for continuous, frequency ranges defined based on Fig. 1e), and the analysis focuses on the top five channels contributing most to the PLS predictions (see Methods Decoder interpretation). LMP shows low PLV in the sustained task but PLV values comparable to other features in the continuous task, suggesting that it follows well the dynamics of faster repetitive flexions but fails to track slower, more static sustained movements. Beta likewise shows reduced PLV during the continuous task. Low-gamma shows PLV values comparable to high-gamma in the sustained task but low PLV during continuous movements. All of these patterns mirror the decoding results. In contrast, high-gamma exhibits the strongest PLV in both tasks, aligning with its superior decoding performance and robustness during generalization.

Based on these results, we focus the remainder of the study on high-gamma activity.

### Short temporal windows enhance generalization

We next examine the decoder’s lookback window to assess how the length of past neural activity used for decoding influences generalization to the unseen condition. Figure 3a shows regression performance for window lengths ranging from 1 s, the duration commonly used in ECoG de-coding ^32^ and the one used in Figure 2, down to 50 ms in 50 ms steps. In the seen condition, shorter windows reduce performance for both tasks, particularly for the sustained task. In contrast, in the unseen condition, shorter windows reduce performance only when models trained on the continuous task are transferred to predict sustained movements (cont2sust), and even in this case the decrease is less pronounced than in the seen condition. For the opposite direction (sust2cont), shorter windows increase performance. Overall, shorter windows yield improved generalization (Fig. 3b), with window lengths below ≈ 250 ms exhibiting significantly smaller performance drops than that of the 1 s baseline.

**Fig. 3.**
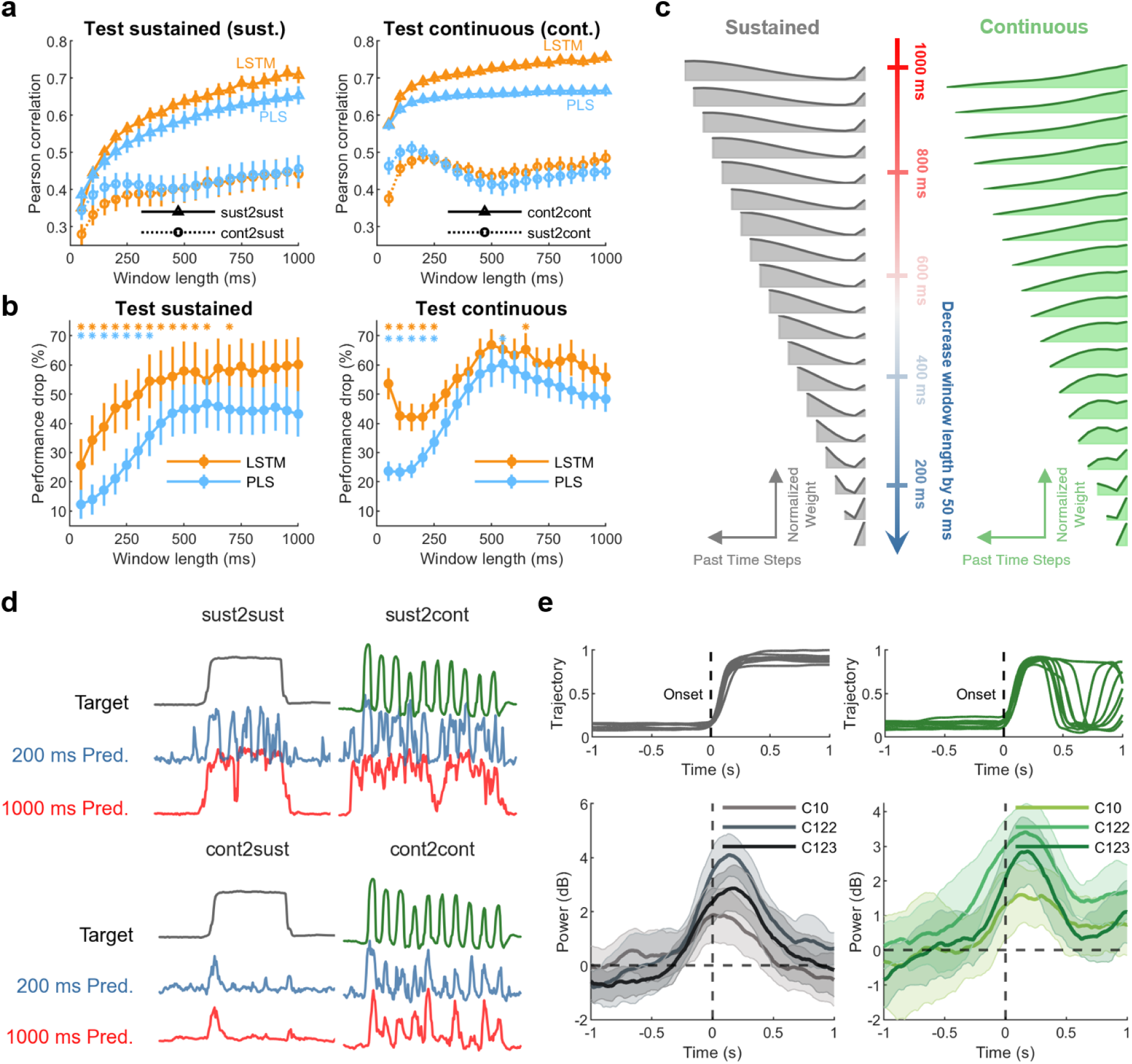
Short temporal windows enhance generalization. **a.** Pearson correlation between dataglove trajectories and model predictions for varying window lengths of past neural activity (50-1000 ms, in 50 ms steps). Both decoders are shown (orange: LSTM; blue: PLS), using high-gamma features in all cases. For the seen condition, mean values (± SEM) are shown with triangular markers and solid lines; for the unseen condition, circular markers and dotted lines. Each point represents 20 values (4 subjects x 5 fingers). **b.** Performance drop in the unseen condition relative to the seen condition for each window length (mean ± SEM). Window lengths with a significantly different performance drop compared to the 1 s window are indicated (\**p <* 0.05, Wilcoxon paired test, FDR-corrected). **c.** Decoder weights assigned to each time step within the window for LSTM models (see Supplementary Fig. S3 for PLS). Left: models trained on the sustained task (grey). Right: models trained on the continuous task (green). Within each window, the leftmost taps represent earlier time points, and the rightmost taps represent recent time points. Larger weights indicate greater contribution to the model’s prediction. **d.** Dataglove trajectories and corresponding LSTM predictions of an example trial (subject 2, thumb) for short (200 ms, blue) and long (1000 ms, red) windows, shown for both the test sustained (left) and test continuous (right) conditions. **e.** Example dataglove trajectories (top) during thumb movements and the corresponding mean high-gamma power (bottom, ± 95% confidence interval) across three ECoG channels from subject 2 (sustained: grey; continuous: green). Trajectories and high-gamma power are aligned to movement onset, with baseline defined using activity from -2 to -1.5 s.

To understand how temporal information is used by the decoders, we examine the contribution of individual time steps within each window (Fig. 3c for LSTM; Supplementary Fig. S3a for PLS). Both models distribute weights across the entire window, but the sustained task model places greater emphasis on earlier time samples, whereas the continuous task model weights more the recent samples. This difference explains why reducing the window length affects sustained-task decoding more than continuous-task decoding in the seen condition (Fig. 3a, solid lines), leading to larger performance decreases, and why shorter windows improve performance when sustained-task models are applied to continuous movements (Fig. 3a, dashed lines, right).

Longer windows limit generalization because they allow models to learn task specific temporal structure rather than features that directly relate to movement. In the sustained task, finger pose remains relatively stable over long periods, encouraging models to rely on slow temporal dependencies (e.g., a flexed finger is likely to remain flexed), which do not transfer to continuous movements (Fig. 3d, sust2cont). Short windows reduce the influence of such task specific structure and force the decoders to rely on the most recent neural activity, which more directly reflects the sensorimotor representation of movement and is shared across tasks.

Findings from movement neurophysiology support this principle^40,41^: high gamma activity linked to movement emerges no more than a few hundred milliseconds prior to movement onset, suggesting that windows longer than this emphasize task level information over muscle related control signals. Our data show a similar pattern, with high gamma increases preceding movement onset by only a few hundred milliseconds (Fig. 3e).

Based on these results, we use a 200 ms window for all subsequent analyses.

### Generalization performance of linear and nonlinear decoders

We examine decoder architecture differences using a 200 ms window. In the seen condition, LSTM achieves significantly better performance than PLS for both tasks (Fig. 4a). However, this advantage disappears in the unseen condition, with no significant difference between the two models, and with PLS outperforming LSTM in cont2sust. The same pattern is observed when performance is assessed using mean-squared error (Fig. 4b). These results, consistent with previous findings from microelectrode studies in non human primates^42^, indicate that the nonlinear model overfits the training data, limiting its ability to generalize across tasks. Example predicted trajectories for both models illustrate this pattern (Fig. 4c).

**Fig. 4.**
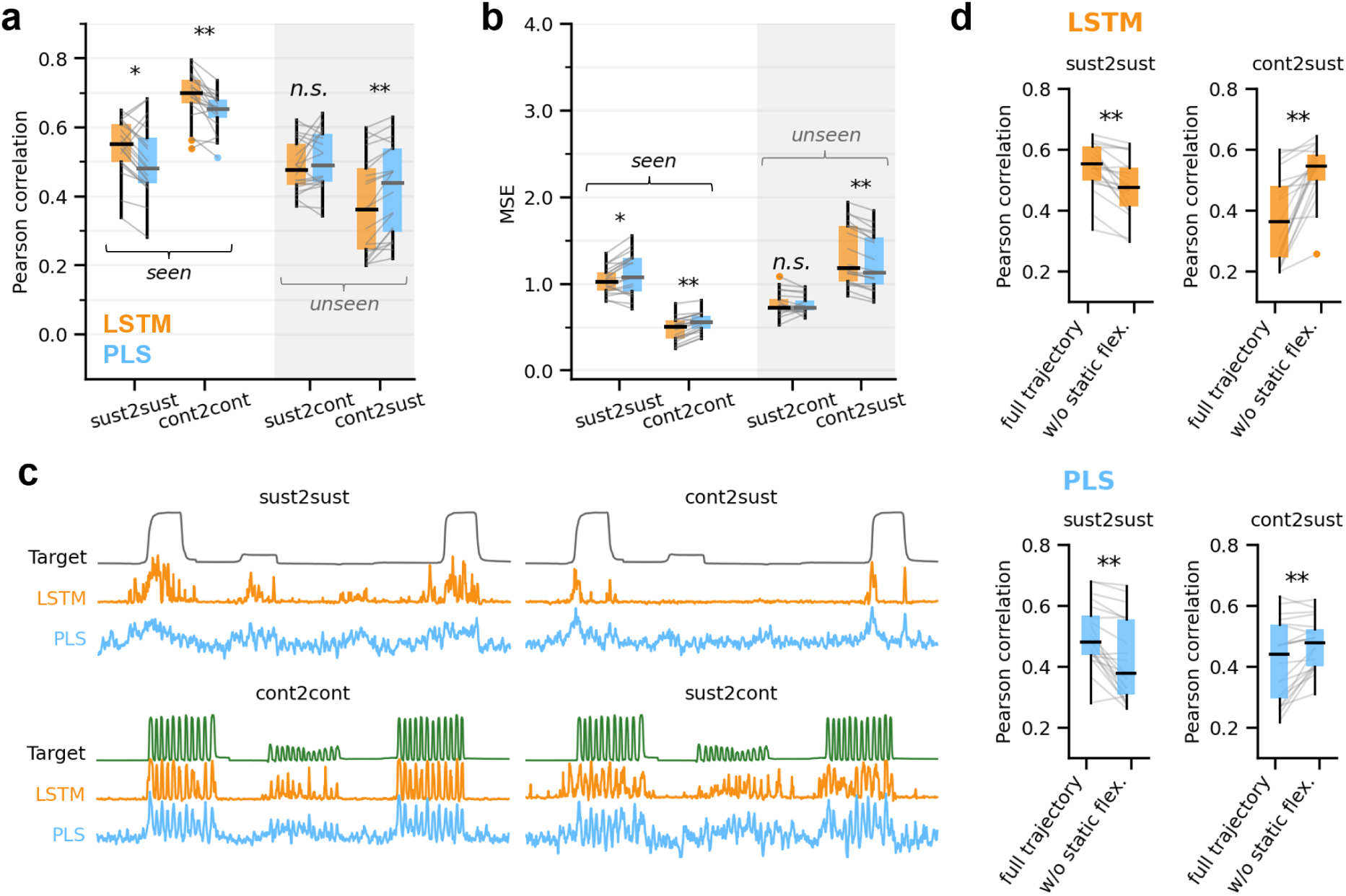
Decoder performance on seen vs unseen tasks and the failure to generalize static flexion. **a.** Pearson correlation between target and predicted trajectories for LSTM and PLS in the seen (left) and unseen (right, grey background) conditions. High-gamma features and 200 ms windows are used. **b.** Same as panel a, but using MSE as the performance metric. **c.** Example dataglove trajectories (target) and corresponding predictions for each decoder in the seen (left) and unseen (right) conditions (subject 1, middle finger). **d.** Effect of removing static flexion samples from the test set on decoding performance. In panels a, b, and d, boxes show the median and IQR range, whiskers show the most extreme non-outlier values (within 1.5 × IQR), and outliers are plotted as dots. Grey lines connect paired samples. Statistical significance is assessed with the Wilcoxon signed-rank test (\**p <* 0.05, \*\**p <* 0.005; FDR corrected).

The trajectories in Fig. 4c also reveal that models trained on the continuous task fail to predict the static flexion samples of the sustained task. This effect is consistent across subjects, suggesting that static poses require explicit representation in the training data. To quantify this, we recompute performance metrics after removing static flexion samples from the test set (Fig. 4d). In the seen condition (sust2sust), removing static-flexion samples (see Fig. 5c) reduces performance, indicating that both models (LSTM and PLS) decode static poses effectively when they are present in the training set. In contrast, in the unseen condition (cont2sust), removing static samples increases performance, demonstrating that these samples are not predicted correctly when models are trained exclusively on dynamic flexion data.

**Fig. 5.**
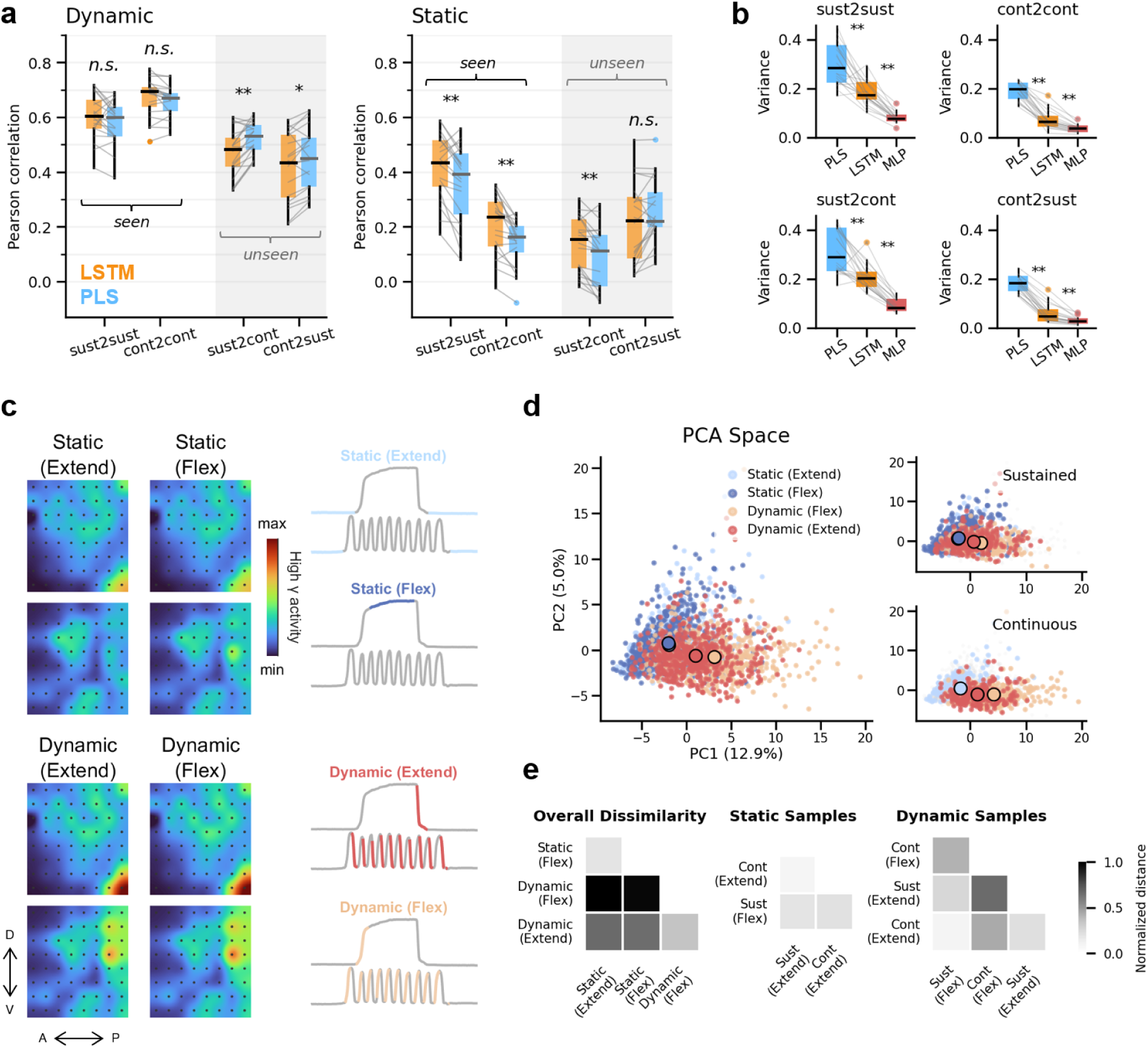
Neural and decoding differences between static and dynamic actions. **a.** Pearson correlation between target and predicted trajectories when only static samples (right plot) or only dynamic samples (left plot) are evaluated, comparing LSTM and PLS in the seen (left boxes) and unseen (right boxes, grey background) conditions. **b.** Variance of the predicted trajectories during static extension for PLS, LSTM, and MLP. Lower variance indicates more stable predictions. **c.** Spatial distribution of high-gamma activity across the two ECoG grids for each action type in subject 3 (thumb movements). Example dataglove traces for each condition are shown to the right. **d.** Dimensionality-reduction of all samples considered in panel c using PCA. Each point represents a high-gamma time sample, colored according to action type. Colored circles indicate the centroid of each action’s cluster. The right panels show the same PCA space when considering only samples from the sustained task or only samples from the continuous task. **e.** Dissimilarity between cluster centroids for the four action types, averaged across all subjects and fingers (n=20), for the PCA space from panel d. The right panels show centroid dissimilarity when clusters from the two tasks are compared separately. In panels a and b, boxes show the median and IQR range, whiskers show the most extreme non-outlier values (within 1.5 × IQR), and outliers are plotted as dots. Grey lines connect paired samples. Statistical significance is assessed with the Wilcoxon signed-rank test (\**p <* 0.05, \*\**p <* 0.005; FDR corrected).

### Neural and decoding differences between static and dynamic actions

#### Linear decoders generalize better to dynamic movements, whereas nonlinear decoders are more stable during static poses

We compare decoding performance for static and dynamic actions separately (Fig. 5a). Notably, the two decoder types exhibit distinct behavior across these actions. PLS shows significantly better performance than LSTM when generalizing dynamic movements across tasks (unseen conditions). In contrast, this advantage disappears for static poses: LSTM outperforms PLS in the sust2cont condition, and the two models do not differ significantly in cont2sust.

Based on the example trajectories in Fig. 4c, we hypothesize that LSTM’s better performance on static movements arises from producing more stable (less noisy) predictions during static periods. To test this, we compute the variance of each decoder’s output during static extension, where the data glove trajectory remains constant and variance reflects prediction noise. LSTM shows significantly lower variance than PLS across all conditions (Fig. 5b). To confirm that this difference comes from model nonlinearity rather than recurrent memory, we evaluate a multilayer perceptron (MLP), a nonlinear model without recurrence, and find that it also produces low variance predictions.

#### Neural activity distinguishes better static vs dynamic actions than flexion vs ex-tension

To understand why static and dynamic actions differ in decodability, we examine the underlying neural activity. We categorize each sample as static extension, static flexion, dynamic extension, or dynamic flexion, pooling across both tasks. Figure 5c shows an example of average high-gamma activity for these four conditions. Static extension and static flexion show strong spatial similarity, and the two dynamic conditions likewise exhibit similar patterns.

To quantify these relationships across subjects and fingers, we apply both PCA and UMAP to the high-gamma features (Fig. 5d left; Supplementary Fig. S6). PCA provides a linear dimensionality-reduction approach, whereas UMAP serves as a nonlinear alternative. In both representations, static and dynamic samples form more distinct clusters than flexion and extension, with both methods showing consistently larger separability between static and dynamic states (Fig. 5e left; Supplementary Fig. S6b, left). Notably, in Fig. 5b all three decoders show higher prediction variance when trained on the sustained task compared to the continuous task. This effect is likely driven by the strong neural similarity between static extension and static flexion. Although these states are neurally alike, they correspond to opposing behavioral outputs (dataglove trajectory values), making it more difficult for sustained-trained decoders to distinguish between them and results in noisier predictions.

#### Static-dynamic structure is preserved across movement tasks

The static-dynamic structure is maintained when considering only sustained task samples or only continuous task samples (Fig. 5d, right), indicating that both tasks share a common underlying neural representation of high-gamma activity. Quantifying inter cluster distances confirms this: clusters from the two tasks lie closer to each other than static and dynamic clusters overall (Fig. 5e, right). Equivalent results were obtained in the UMAP space (Supplementary Fig. S5). Collectively, these findings suggest that BCI generalization may benefit from approaches that explicitly model static versus dynamic actions, as this distinction is robust and preserved across tasks.

### Nonlinear decoders benefit more from diverse training sets

Prior work with microelectrode recordings has shown that neural networks generalize better to multiple tasks when the training set includes examples from all those tasks, mitigating overfit-ting ^42^. We test whether the same principle applies to high density ECoG (Fig. 6). Starting from the unseen case (0% seen), we progressively add samples from the target task to the training set up to 50%, and evaluate decoding performance for the full trajectory as well as for its dynamic and static components.

**Fig. 6.**
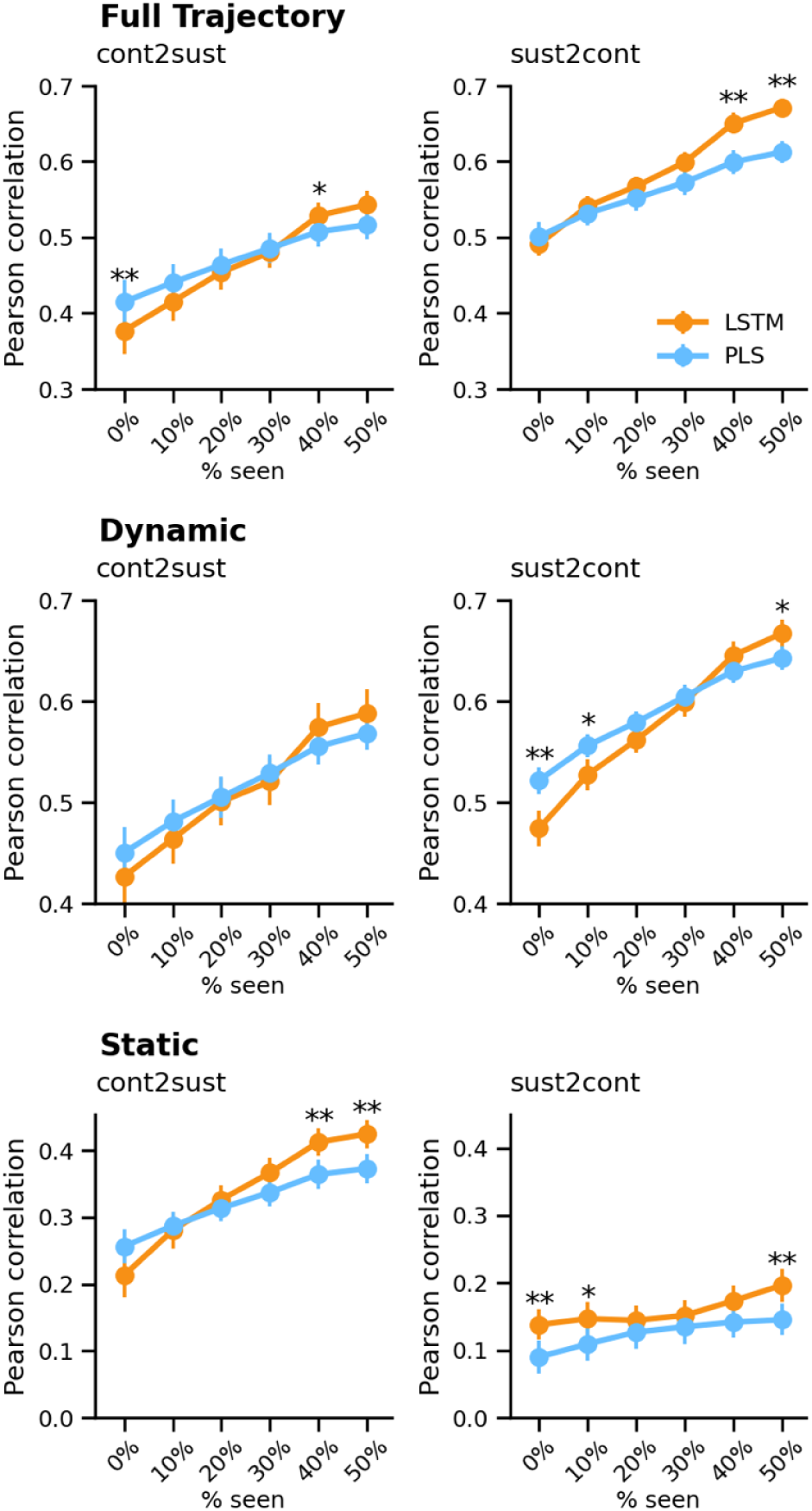
Nonlinear decoders outperform when both tasks are represented in the training set. Pearson correlation (mean ± SEM) between dataglove trajectories and model predictions for increasing percentages of target-task data included in the training set, ranging from 0% (unseen case) to 50% (equal representation of sustained and continuous tasks). Results are shown for both decoders (orange: LSTM; blue: PLS) when considering the full trajectory (top), dynamic samples only (middle), and static samples only (bottom). Significant differences between decoders are assessed using the Wilcoxon signed-rank test (\**p <* 0.05; \*\**p <* 0.005; FDR corrected).

Performance increases steadily for both LSTM and PLS as more samples from the target task are included in training. This improvement is expected, as exposure to the target task reduces the generalization gap and increases the total amount of training data. However, the gain is consistently stronger for LSTM than for PLS. As a result, LSTM begins to significantly outperform PLS in settings where no difference is observed in the unseen case (e.g., sust2cont for full trajectories and cont2sust for static samples), and even in cases where PLS originally outperforms LSTM (e.g., cont2sust full trajectories and sust2cont dynamic samples). These results show that, similar to microelectrode studies, nonlinear decoders trained on ECoG signals benefit substantially from diverse training sets, and their generalization improves more rapidly than that of linear models when both tasks are represented.

### Anatomical differences influence decoder generalization

Lastly, we examine how anatomical location influences generalization performance. We first quantify the contribution of each ECoG channel to the decoder to assess whether motor or sensory regions are more informative for each task (Fig. 7a,b for PLS; Supplementary Fig. S7 for LSTM). Across subjects, both motor and sensory channels contribute to decoding, but the relative contribution of each region varies by subject, likely reflecting differences in grid placement. Subjects with greater motor coverage (e.g., subjects 2 and 4) show correspondingly larger motor contributions. Overall, sensory channels tend to contribute more strongly during the continuous task than during the sustained task.

**Fig. 7.**
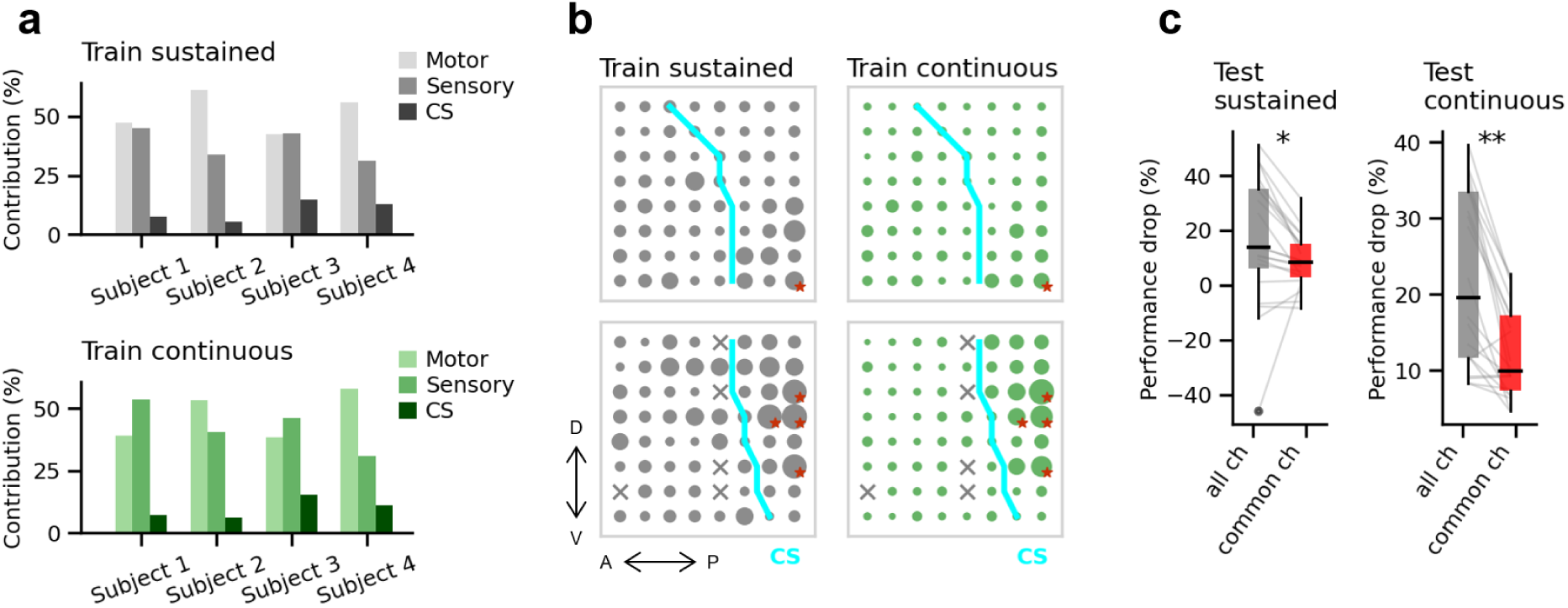
Anatomical differences influence decoder generalization. **a.** Contribution of each anatomical region (motor: anterior to the central sulcus; sensory: posterior to the central sulcus; CS: channels directly over the sulcus) to the PLS decoder for each subject and each training task. **b.** Channel-wise contributions to the decoder for the two ECoG grids of subject 1. Larger circles indicate stronger contributions. Channels rejected during preprocessing are marked with an “x”. The central sulcus is shown in cyan. The top five channels contributing most strongly across both tasks are marked with red stars (see Supplementary Fig. S7 for all subjects and LSTM results). **c.** Performance drop in the unseen condition when using either all channels or only the five common channels identified in panel b. Boxes show the median and IQR range, whiskers show the most extreme non-outlier values (within 1.5 × IQR), and outliers are plotted as dots. Grey lines connect paired samples. Statistical significance is assessed with the Wilcoxon signed-rank test (\**p <* 0.05, \*\**p <* 0.005; FDR corrected).

Figure 7b shows an example of channel wise contributions for both training tasks. Channels anterior and posterior to the central sulcus are categorized as motor and sensory, respectively, and channels directly over the sulcus are labeled CS. Channel contributions differ between the two tasks, an effect that is consistent across subjects and aligns with prior suggestions that static and dynamic movements rely on partially distinct sensorimotor subregions.

We next test whether anatomical specificity contributes to the generalization drop observed across tasks. If performance loss arises from task-dependent channel selectivity, then restricting decoding to channels that are informative for both tasks should improve generalization. To evaluate this, we select during PLS training (with the test set kept separate) the top five channels with the highest combined contribution across sustained and continuous tasks. Using only these common channels reduces the performance drop in the unseen condition compared with using all channels (Fig. 7c), indicating improved generalization. Interestingly, examining the anatomical locations of these common channels (Supplementary Fig. S8) shows that they lay predominantly in the sensory cortex, suggesting that sensory representations generalize more reliably across tasks than motor representations.

## Discussion

Our findings show that generalization across static and dynamic actions is shaped by multiple components of the BCI decoding pipeline, indicating that design choices optimized for a specific motor task will not necessarily promote generalization. This has important implications for the development of motor BCIs, particularly those targeting fine finger movements which rely on a diverse set of actions supported by distinct neural representations. Future BCI systems will therefore need to incorporate decoding strategies that adapt across tasks and movement types rather than depend on features or architectures tuned to a single scenario.

Identifying high-gamma activity as the optimal feature for cross-task generalization aligns with extensive prior evidence that links high-gamma power to movement-related cortical activation^38^. Our results extend this evidence by demonstrating that high-gamma features also support continuous prediction of previously unseen motor tasks across different movement types. In addition to high-gamma, LMP has been widely used in intracortical motor BCIs^43^. In our data, LMP performed well only during the highly dynamic task (continuous task) and did not generalize to the sustained task, with more static postural components. This does not contradict earlier studies reporting robust LMP decoding ^5,14^, as those experiments typically focused on dynamic behaviors. Nevertheless, some reports have described LMP-based decoding of slower or more static postures ^28,44^. A possible explanation for this discrepancy is that static postures in those studies were brief (1–2 s), whereas in our paradigm the posture was maintained for 4 s. In addition, those studies used standard (i.e. non-high-density) ECoG arrays, which aggregate LMP over a broader cortical region and may yield different spatial-temporal signal properties. Turning to low-gamma activity, previous work has shown that this band shows sustained activity during prolonged static poses ^23,24,26,45^, suggesting that it may provide a useful feature for decoding maintained postures. Our results are consistent with this observation: low-gamma features performed significantly better in the sustained task (Fig. 2c). Nonetheless, high-gamma remained the most effective feature overall, and adding low-gamma did not yield any significant improvement in decoding performance (Supplementary Fig. S2).

A common practice in ECoG-based BCIs is to use relatively long windows of recent neural activity, typically around 1 s, for decoding. This approach is used not only in studies aimed at optimizing decoding pipelines ^11,17^, but also in applied research developing assistive strategies for individuals with paralysis ^32,46^. Such window lengths have generally performed well in these contexts, likely because the decoding targets in those studies predominantly involve dynamic arm and hand movements rather than static postures. Our results indicate that when both dynamic and static actions are relevant, shorter temporal windows (<250 ms; Fig. 3) provide better generalization across tasks. In contrast to the ECoG literature, the use of short decoding windows is already common in microelectrode-based BCIs^8,47–49^. Although these studies rely on averaged firing rates rather than high-gamma power, the two measures have been shown to correlate strongly ^50,51^, suggesting that similar temporal principles may apply across recording modalities.

The advantage of shorter windows for enhancing generalization is consistent with the temporal dynamics of high-gamma activity, which precedes movement onset by only a few hundred milliseconds in primary sensorimotor cortex ^40,41^. Our findings therefore apply most directly to BCIs that rely on signals from this region, which remains the most commonly targeted region in the field due to its strong involvement in movement execution and sensory feedback^52^. Recent studies, however, have begun to investigate whether incorporating additional cortical areas could improve BCI performance ^53–55^. Some of these regions, such as premotor and posterior parietal cortices, show intention-related activity that can precede movement onset by up to a few seconds ^56^. In such cases, longer temporal windows might be beneficial for capturing this anticipatory information. Further research is required to determine how decoding strategies should adapt when BCIs integrate these additional brain regions, and to clarify how temporal window selection interacts with the specific neural processes each area encodes.

Another consideration for generalization is decoder architecture. Recent BCI studies have successfully employed both linear ^49,57^ and nonlinear decoders ^8,46,47^, as well as hybrid approaches ^32,48,58^, and there is currently no consensus on which architecture is preferable. Our results suggest that the optimal decoder choice depends on the context in which the BCI is developed. When the goal is to optimize performance within a specific task, or when substantial training data is available, nonlinear decoders tend to perform best. In contrast, when generalization across tasks is required and only limited single-subject data can be collected, linear decoders offer more robust performance, particularly for highly dynamic tasks.

Regarding action types, our results indicate that both static and dynamic actions must be represented in the training data to enable generalization. When one action was absent from the training set, in our case static flexion was missing from the continuous task, decoders struggled to predict it (Fig. 4d). This outcome aligns with neurophysiology studies showing that static and dynamic actions rely on partly distinct neural mechanisms ^21–24^. More broadly, our findings reveal a stable static-dynamic neural structure that is preserved across both tasks, suggesting that BCI generalization may benefit from approaches that explicitly model these two action types. One potential strategy is velocity decoding. Static postures naturally correspond to zero velocity, whereas dynamic movements correspond to positive or negative velocities for flexion and extension, respectively. However, in our case, decoding velocity did not improve performance (Supplementary Fig. S4), likely because flexion and extension show very similar neural representations (Fig. 5d). When only velocity magnitude (i.e. speed) is predicted, which removes directional information, performance increases. This is consistent with known limitations of ECoG recordings for distinguishing movement direction^20^.

Another strategy is to use a hierarchical or “state based” decoding framework, in which an initial classifier discriminates between static and dynamic states, followed by specialized decoders that resolve the specific action (e.g., flexion versus extension). Although state based models have been proposed in the BCI literature ^32,35,58^, to our knowledge they have only been explicitly applied to static versus dynamic differentiation in one preliminary study ^59^. In our dataset, hierarchical decoders outperform unified models only for the continuous task (Supplementary Fig. S5), whereas unified models remain superior for the sustained task. The limited advantage of hierarchical decoders may reflect dataset limitations: training multiple specialized decoders divides the available training samples across models, and our dataset (∼15 minutes per subject) is insufficient to fully support both levels of the hierarchy. Studies with substantially larger datasets will be needed to assess the full potential of static-dynamic state modeling for BCI generalization. Overall, although the preserved static and dynamic structure suggests promising possibilities for improving generalization across tasks, further work is needed to determine how this structure can be best used within practical BCI applications, and to assess how well it is maintained in individuals suffering from paralysis^60^.

From an anatomical perspective, our results show that decoders find more shared contributions across tasks in somatosensory regions than in motor regions, and that focusing on these shared regions enhances generalization. Somatosensory cortex is not an uncommon target for BCIs: prior studies have demonstrated high decoding accuracy from this area ^61,62^, even in individuals without sensory feedback, as it preserves neural responses to attempted movements years after paralysis ^63^. Overall, our findings suggest that electrode-placement strategies for motor BCIs, often guided by task-related fMRI activations^64^, should consider the full range of intended tasks. This includes accounting not only for the relevant body parts but also for both static and dynamic actions ^18^, which may rely on different cortical representations.

Our study focuses on finger movements, given their complexity and relevance for many daily activities. However, similar distinctions between static and dynamic actions have been reported for other muscle groups, including the hand ^23^, the arm ^65^, and even the small facial muscles involved in speech ^66^. These similarities suggest that the principles identified here may extend to other BCI scenarios requiring generalization across tasks.

It is important to note that our findings apply to BCIs that aim to generalize across tasks. In some scenarios generalization is not desirable, as BCIs are designed around a single, well defined application, where exploiting task structure is advantageous to enhance performance. Examples include BCIs that rely on language models ^67^ or AI assisted motor control^68^ approaches to learn contextual information to complete the intended action. In such cases, our conclusions regarding generalization do not apply. Ultimately, the appropriate design strategy depends on the specific goals of the BCI and whether flexibility across tasks is required.

Finally, this study has several limitations that should be considered. Because ECoG recordings are only available in infrequent clinical circumstances, our sample size is necessarily small. Nevertheless, having four subjects with comparable high-density ECoG coverage of the sensorimotor cortex provides a valuable dataset, and the significance of effects supports the robustness of our findings. Data collection was also constrained by the brief intraoperative time window, yielding approximately 15 minutes of recordings per subject. While this was sufficient for the decoders evaluated here, it limited our ability to explore alternatives, such as state based decoders, and restricted the number of tasks we could include. We selected two tasks that capture the principal types of dynamic and static finger actions, which should support generalization to additional tasks that draw on combinations of these actions. Even so, future work will be needed to test how the principles identified here extend to additional motor tasks.

In summary, our findings reveal that generalization across static and dynamic actions depends on multiple components of the BCI pipeline, including feature choice, lookback window, decoder architecture, and anatomical location. These insights highlight the importance of explicitly considering task diversity when designing BCIs intended to operate across multiple movement scenarios. Looking forward, future work should explore whether the principles identified here extend to additional tasks and muscle groups, more complex behaviors, and broader cortical regions, as well as evaluate their implications for BCI systems intended for real-world use.

## Methods

### Participants

Five patients (Four males; age range: 30-60 years) undergoing glioma surgery were recruited from the Department of Neurosurgery, West China Hospital, Sichuan University. To maintain consistency across neural recordings, only patients undergoing left-sided awake craniotomies were included. All participants performed right-hand movement tasks. Study inclusion required: (1) preserved motor function with the tumor sparing the primary hand sensorimotor cortex as confirmed by preoperative MRI; (2) absence of psychiatric illness, neurological comorbidities, or substance abuse; and (3) the capacity to cooperate with intraoperative behavioral tasks. One patient (male) was subsequently excluded from the analysis due to suboptimal task compliance during the awake phase. The remaining four participants were designated as subjects 1-4.

The study protocol received approval from the Institutional Review Board of West China Hospital (No. 20242138) and adhered to the Declaration of Helsinki. Written informed consent was obtained from all participants. Data sharing and management were conducted under the Data Processing Agreement between KU Leuven and West China Hospital, effective as of May 1, 2024.

### Intraoperative mapping and ECoG acquisition

Tumor resection was performed via awake craniotomy to maximize safe resection while safeguarding functional integrity. Following craniotomy, the lesion’s boundaries were initially estimated using a neuronavigation system (BrainLab AG, Munich, Germany). To precisely delineate the hand-specific motor representation in the left hemisphere, functional mapping was performed via bipolar direct cortical stimulation (DCS). Positive motor responses were defined as evoked movements in the contralateral (right) fingers, while sensory responses were identified based on the patient’s self-report of paresthesia. These functional landmarks (DCS-positive sites) were used to identify the precentral and postcentral gyri.

Intraoperative photographs of the exposed cortex were captured, and DCS-positive sites were marked with 3 mm radius digital indicators. Subsequently, two 8 × 8 platinum-electrode ECoG grids (HKHS Healthcare Co., Ltd., Beijing, China) were placed over the identified sensorimotor area. Each electrode featured a 2 mm diameter contact with a 5 mm center-to-center spacing.

Neural signals were recorded using a NeuSen H system (Neuracle Technology Co., Ltd., Changzhou, China) at a sampling rate of 2000 Hz. Ground and reference signals were obtained via needle electrodes placed on the ipsilateral and contralateral scalp, respectively. For spatial verification, the final grid positions were photographed and co-registered with the DCS-labeled cortical maps to ensure precise anatomical-functional correspondence of the recorded electrodes.

### Finger task design

Participants performed two types of finger-movement tasks, sustained and continuous, using the right hand (contralateral to the ECoG grids) during the intraoperative recording sessions. Four distinct movement types were performed: (1) thumb, (2) index finger, (3) combined middle-ring-little (MRL) finger, and (4) full-fist flexion-extension. Finger kinematics (trajectories) were captured at 20 Hz via a 5DT Data Glove 5 Ultra (5DT, Irvine, CA, USA).

Each trial followed a structured sequence (Fig. 1a,b): a fixation phase (1.5 s), a cue phase (1 s) indicating the target finger, a preparation phase (2 s), a task phase (4 s for sustained; 7 s for continuous), and a rest phase (2 s). To minimize visual-evoked confounding effects and precisely control movement timing, we employed a shrinking-circle paradigm^9,69^. For the sustained task, participants were instructed to initiate finger flexion when the circle reached the outer cross-bars and maintain the flexed pose until the circle contacted the inner cross-bars (4 s), after which they returned to the neutral pose. For the continuous task, participants performed repetitive, rhythmic flexion-extension of the cued fingers throughout the 7-second task phase as the circle moved.

The experimental paradigm was implemented in Psychtoolbox-3. Within each experimental block, the presentation order of sustained and continuous tasks was pseudo-randomized. Furthermore, the four finger-movement trials within each task were presented in a pseudo-random sequence. Each participant (subjects 1-4) completed eight blocks, resulting in approximately 15 minutes of total ECoG and finger kinematic recording. All participants had a practice session the day before the experiment session to become familiar with the paradigm.

### ECoG and kinematic data preprocessing

The raw continuous ECoG recording were band-pass filtered with a zero-phase, 4th-order IIR Butterworth filter between 0.15 and 200 Hz. To eliminate power-line interference, 3rd-order notch filters were employed at 50 Hz and its associated harmonics (100 Hz and 150 Hz). Channel integrity was then assessed visually and by calculating the variance of each electrode; noisy channels were rejected and excluded from further analysis. This step resulted in a final dataset of 123, 124, 127, and 127 clean channels for subjects 1-4, respectively. To enhance the signal-to-noise ratio and mitigate common-mode artifacts, the ECoG signals were re-referenced to the common average of all clean channels. Finally, the ECoG signals were downsampled to 1000 Hz to reduce computational complexity while preserving relevant spectral information.

Finger movement trajectories, initially captured at 20 Hz with 12-bit resolution, were synchronized with the ECoG recordings by resampling the data to 1000 Hz. To mitigate edge artifacts, we applied a 5-second mirror padding to both the beginning and the end of each raw trajectory. The padded signals were then resampled using a polyphase antialiasing filter (via MATLAB’s *resample* function). Following this, the temporary padding was removed to restore the original signal duration. This procedure ensured a unified sampling rate and precise temporal alignment between the kinematic trajectories and the ECoG trial markers.

### Movement onset detection and temporal alignment

To investigate movement-related power changes (cf., Fig. 1g and Fig. 3e), individual ECoG trials were aligned to the movement onset (*t* = 0). Finger trajectories were first smoothed using a 100-ms moving average and normalized to a range of [0, 1]. Onset detection was performed using a velocity-based dual-threshold algorithm applied to the representative finger trajectory for each task (thumb for thumb trials; index for index and fist trials, middle for MRL trials). The algorithm first identified a reference point as the first significant peak exceeding 50% of the trial’s maximum velocity, with the velocity profile derived from the first-order derivative (via MATLAB *diff* function) of the movement trajectory. A backward search was then conducted within a 0.5-second window from this peak to locate the first time point where the velocity exceeded 20% of the maximum. This point was defined as the movement onset. All kinematic trajectories and corresponding neural features were then centered at this detected onset, enabling consistent across-trial visualization of sustained and continuous movements, as well as the resulting event-related spectral perturbations.

### Modeling

#### Feature extraction

Neural features were extracted from the 1000 Hz ECoG signals using a multi-band filtering approach. To mitigate edge effects during filtering, we applied a 1-second mirror padding to the signal boundaries, and removed the padded segments once feature extraction was done. Seven distinct features were then derived using zero-phase (two-pass) Butterworth filters and the Hamming window (using FieldTrip toolbox ^70^):

- Local Motor Potential (LMP): Extracted via a 5*^th^*-order low-pass filter at 3 Hz to capture low-frequency temporal fluctuations.
- Spectral Envelopes: We calculated the analytic signal envelope using the Hilbert transform after filtering. The bands included: Delta (*δ*): < 5 Hz (5*^th^*-order low-pass); Theta (*θ*): 5-8 Hz (3*^rd^*-order band-pass); Alpha (*α*): 8-12 Hz (3*^rd^*-order band-pass); Beta (*β*): 12-34 Hz (3*^rd^*-order band-pass); Low gamma (low *γ*): 34-60 Hz (4*^th^*-order band-pass); High gamma (high *γ*): > 60 Hz (7*^th^*-order high-pass).

Finally, both neural features and kinematic data were downsampled to 20 Hz to match the BCI decoder’s prediction rate.

#### Decoders

##### Task formulation

We formulated the finger-movement predition task as a continuous regression problem, where the goal is to map ECoG features to the kinematic trajectories of individual fingers. Let *y*(*t*) ∈ ℝ represent the normalized trajectory of a specific finger (e.g., thumb or index) at time *t*. The input to the decoder consists of multi-channel time-frequency features extracted from a lookback window of length *T* . At each time step *t*, the input feature tensor is defined as: **<u>X</u>***_t_* ∈ ℝ*^C×T^ ^×F^*where: *C* is the number of ECoG channels; *T* is the number of temporal bins in the lookback window; *F* is the number of spectral frequency bands (e.g., LMP, *δ*, *θ*, *α*, *β*, low *γ*, and high *γ*). The decoding objective is to learn a linear/nonlinear mapping function *f* (·) such that the predicted trajectory *y*^(*t*) minimizes the error relative to the ground-truth trajectory *y*(*t*): *y*^(*t*) = *f* (**X***_t_*; *θ*) where *θ* denotes the trainable parameters of the decoder.

##### Linear decoder

We used Partial Least Squares (PLS) regression as a linear decoder due to its robust performance in modeling high-dimensional neural data ^71^, a capability that has underpinned several multi-way extensions in recent ECoG-BCI studies^32,58^. To accommodate the PLS model, the training feature tensors were vectorized into a matrix **X** ∈ ℝ*^N×^*^(*C·T*^ *^·F^* ^)^, where *N* denotes the total number of training samples generated via a sliding window. PLS identifies a set of latent components by simultaneously decomposing **X** and the corresponding kinematic trajectories **y** ∈ ℝ*^N^* . The optimization objective is to maximize the pairwise covariance between these components, ensuring that the extracted latent features capture the most predictive information for finger movement. In this study, the number of latent components was selected from the range [1,50] using cross-validation. Models were fitted using the PLSRegression class from scikit-learn (1.3.0).

##### Nonlinear decoder

We trained a Long Short-Term Memory (LSTM) network as our non-linear decoder, given its demonstrated efficacy in sequential neural modeling ^37^ and its wide application in recent ECoG literature^12,17,33^. The network comprised a single LSTM layer with 256 hidden units. At each time step, the model processed the sequence **X** ∈ ℝ*^T^ ^×^*^(*C·F*)^ from the lookback window, and the hidden state of the final temporal bin was extracted for prediction. To enhance stability and prevent overfitting, this state underwent Layer Normalization ^72^ followed by a Dropout layer (*p* = 0.5). A two-layer Multilayer Perceptron (MLP) (256 → 128 → 1) with ReLU activation was employed to transform the normalized features into the final finger trajectory prediction. To evaluate the impact of recurrent temporal modeling, we further implemented a three-layer MLP as a non-recurrent nonlinear baseline. Unlike the LSTM, this model received vectorized input **X** ∈ ℝ*^C·T^ ^·F^* and processed it through a sequence of linear layers (*C* · *T* · *F* → 256 → 128 → 1). Each hidden layer utilized ReLU activation and Dropout (*p* = 0.5) to mitigate overfitting.

To train the neural networks, we used the Adam optimizer with a constant learning rate of 10*^−^*^3^ and a mean squared error (MSE) loss function. The training process is governed by an early stopping criterion with a patience of 10 epochs. Training is conducted for a maximum of 100 epochs with a batch size of 128.

#### Decoder training and evaluation

##### Chronological cross-validation and data splitting

To ensure a robust assessment of decoder performance while respecting the temporal structure of neural data, we implemented a five-fold chronological cross-validation scheme. The continuous ECoG feature envelopes and corresponding trajectories were divided into five equal temporal segments. In each fold, one segment was reserved for testing, one for validation (hyperparameter tuning and early stopping), and the remaining three were concatenated for training. This split was synchronized across neural and kinematic modalities. To prevent data leakage, normalizations (z-score standardization) was performed using parameters (mean and standard deviation) estimated solely from the training set and then applied to the validation and test sets.

##### Seen and unseen conditions

To evaluate the decoder’s generalizability across different movement tasks, we conducted both seen and unseen experiments based on trial markers:

- Seen condition: Decoders were trained and tested on the same task type, yielding **sust2sust** and **cont2cont**.
- Unseen condition: Decoders were trained on one movement type (e.g., sustained) and evaluated on the other (e.g., continuous), yielding **sust2cont** and **cont2sust**.

##### Sample generation and windowing

Training and testing samples were generated using a sliding window approach with a step size of 1 (corresponding to 50 ms at 20 Hz). At each time step, the input **X** ∈ ℝ*^C×T^ ^×F^* was constructed using a lookback window of *T* temporal bins. Initially we used a 1-second window (*T* = 20), following prior ECoG studies. However, after analysing different window lengths, we adopted a shorter window (200 ms, *T* = 4), which improved generalization performance.

##### Performance metric

We quantified decoding performance using the Pearson correlation between the predicted and ground-truth trajectories for each finger^5^. This metric assesses the linear relationship and temporal synchrony between the decoded kinematics and the actual finger movements. In addition, we computed the mean squared error (MSE), which complements correlation by evaluating the absolute magnitude of prediction errors.

#### Decoder interpretation

We looked into the models’ behavior to identify which temporal bins within the lookback window and which ECoG channels most significantly contributed to the movement prediction. This analysis was restricted to the seen conditions (sust2sust and cont2cont) using high-gamma envelope sequences as inputs.

For PLS, we examined the coefficients assigned by the model to each input feature and grouped them either across temporal bins (Fig. S3a) or across ECoG channels (Fig. 7b and S7). Because features were normalized prior to decoding, the resulting coefficients can be interpreted as indicators of each feature’s contribution to the decoding performance. To allow combining data across participants, we normalized the coefficients separately for each finger and subject.

For LSTM, we employed Integrated Gradient (IG)^73^, an axiomatic attribution method to interpret deep networks. We generated a distribution of 10 baselines by calculating the mean of each ECoG channel across the training set and augmenting it with Gaussian noise. For each input test sample (sized *T* ×*C*), IG attributions were calculated relative to each of these noisy baselines and subsequently averaged to ensure stable and unbiased estimates. The resulting attribution tensors, matching the input dimensions, were aggregated to yield distinct importance profiles. For temporal importance, we calculated the absolute sum of attributions across all ECoG channels for each time step to visualize the model’s reliance on historical neural activity. For spatial importance, we summed the absolute attributions over the temporal dimension to identify the most contributory ECoG channels, providing a spatial topography of movement-related neural features.

To detect channels with high contribution across both sustained and continuous tasks, we extracted the channels’ contribution for each subject, finger, and cross-validation fold and ranked channels by their aggregated weight in each task. For every combination of subject, finger, and fold, we then combined the two task-specific rankings and selected the five channels with the smallest joint rank, yielding the channels most consistently weighted in both conditions.

#### Phase Locking Value

We computed phase-locking values (PLVs) between data glove trajectories and neural features in two frequency ranges: < 0.5 Hz for the sustained task and 0.5-3 Hz for the continuous task. All signals were first band-pass filtered in the frequency range of interest using a zero-phase, 4th-order IIR Butterworth filter. We then obtained the instantaneous phase by taking the angle of the analytic signal derived from the Hilbert transform. For each channel, the PLV was computed as in Lachaux et al. ^39^, reflecting the consistency of phase differences over time.

### Static and dynamic action analysis

#### Action segmentation

We segmented finger movements into dynamic and static actions based on trajectory velocity and position (cf., Fig. 5c). The resulting segments were used to extract time-aligned high-gamma envelopes across all ECoG channels.

##### Dynamic action

The instantaneous velocity was calculated as the first-order derivative (via MATLAB *diff* function) of the finger trajectory. Dynamic actions, both flexion and extension, were identified when the absolute velocity exceeded 10% of its peak value within each task. To ensure temporal continuity, a moving average filter (window size n = 15 for sustained; n = 2 for continuous) was applied to the resulting binary masks.

##### Static action

Periods with velocity below the threshold were categorized as static. Static actions were further subdivided into static-flexion and static-extension. Static-flexion was determined if the amplitude of trajectory exceeded the *mean* + 0.5 × *interquartile range* (*IQR*); otherwise, it was labeled as static-extension. In the continuous task, as only a few samples were labeled as static-flexion, we did not include them for visualization in Fig. 5d-e.

#### Neural manifold analysis

We employed manifold learning to investigate the latent structure of high-gamma activity across dynamic and static actions. To ensure unbiased representation, we first generated a balanced dataset by sub-sampling neural features (channel-wise high-gamma envelopes) across all subjects, fingers, and tasks for four distinct actions: static extension, static flexion, dynamic extension, and dynamic flexion. Dimensionality reduction was performed using both linear Principal Component Analysis (PCA) and nonlinear Uniform Manifold Approximation and Projection (UMAP) ^74^. Neural features were projected onto a two-dimensional manifold space to visualize action-specific clusters. For UMAP, hyperparameters were set to 50 neighbors and a minimum distance of 0.1.

To quantify the separability of neural states, we calculated the Euclidean distance to the centroid for each action cluster within the manifold space. Dissimilarity matrices were then constructed to evaluate the overall distance between the four motor states. Furthermore, we performed subset analyses to compare action representations across the sustained and continuous tasks. All distance metrics were normalized to the global maximum dissimilarity to facilitate cross-condition comparisons.

#### Statistics

Statistical comparisons were performed using the two-sided Wilcoxon signed-rank test for paired samples, unless otherwise specified. The threshold for statistical significance was set at *α* = 0.05. False Discovery Rate (FDR) correction was applied in multiple comparisons. Statistical significance is denoted as follows: \**p <* 0.05, \*\**p <* 0.005. All data are presented as mean ± standard error of the mean (SEM) unless otherwise stated.

## Data availability

Behavioral and electrocorticographic data that support the findings of this study will be deposited in a public repository (OpenNeuro) upon publication.

## Code availability

Code used for experimental analyses, model training, and visualization will be made publicly available upon publication in Zenodo at: https://doi.org/10.5281/zenodo.18887931.

## Acknowledgements

ECM is supported by the Belgian Fund for Scientific Research - Flanders (1102925N). QS is supported by the China Scholarship Council (No.202206050022). YY is supported by funding from National Natural Science Foundation of China (81902532), Sichuan Association for Science and Technology (2025YFRG0006), and the 1.3.5 project for disciplines of excellence-Clinical Research Fund, West China Hospital, Sichuan University (2025HXFH048). JH is supported by the 1.3.5 project for Disciplines of 1435 Excellence Grant from West China Hospital under Grant ZYYC22001 and the Fundamental Research Funds for the Central Universities under Grant YJ202373. MMVH is supported by research grants received from Horizon Europe’s Marie Sklodowska-Curie Action (grant agreement No. 101118964), Horizon 2020 research and innovation programme under grant agreement No. 857375, the special research fund of the KU Leuven (C24/18/098), the Belgian Fund for Scientific Research – Flanders (G0A4321N, G0C1522N, G031426N), and the Hercules Foundation (AKUL 043).

## Author contributions

ECM: Conceptualization, Methodology, Software, Formal analysis, Visualization, Writing-original draft. QS: Methodology, Software, Formal analysis, Visualization, Writing-original draft. YQ, TC, MM, YL, QM, and YY: Resources, Investigation (surgery). YW, JL, and JH: Methodology, Investigation (data collection), Data Curation. All authors: Writing-review & editing. MMVH: Supervision, Project administration, Funding acquisition.

## Competing interests

The authors declare no competing interests.

## Supplementary Figures

**Fig. S1.**
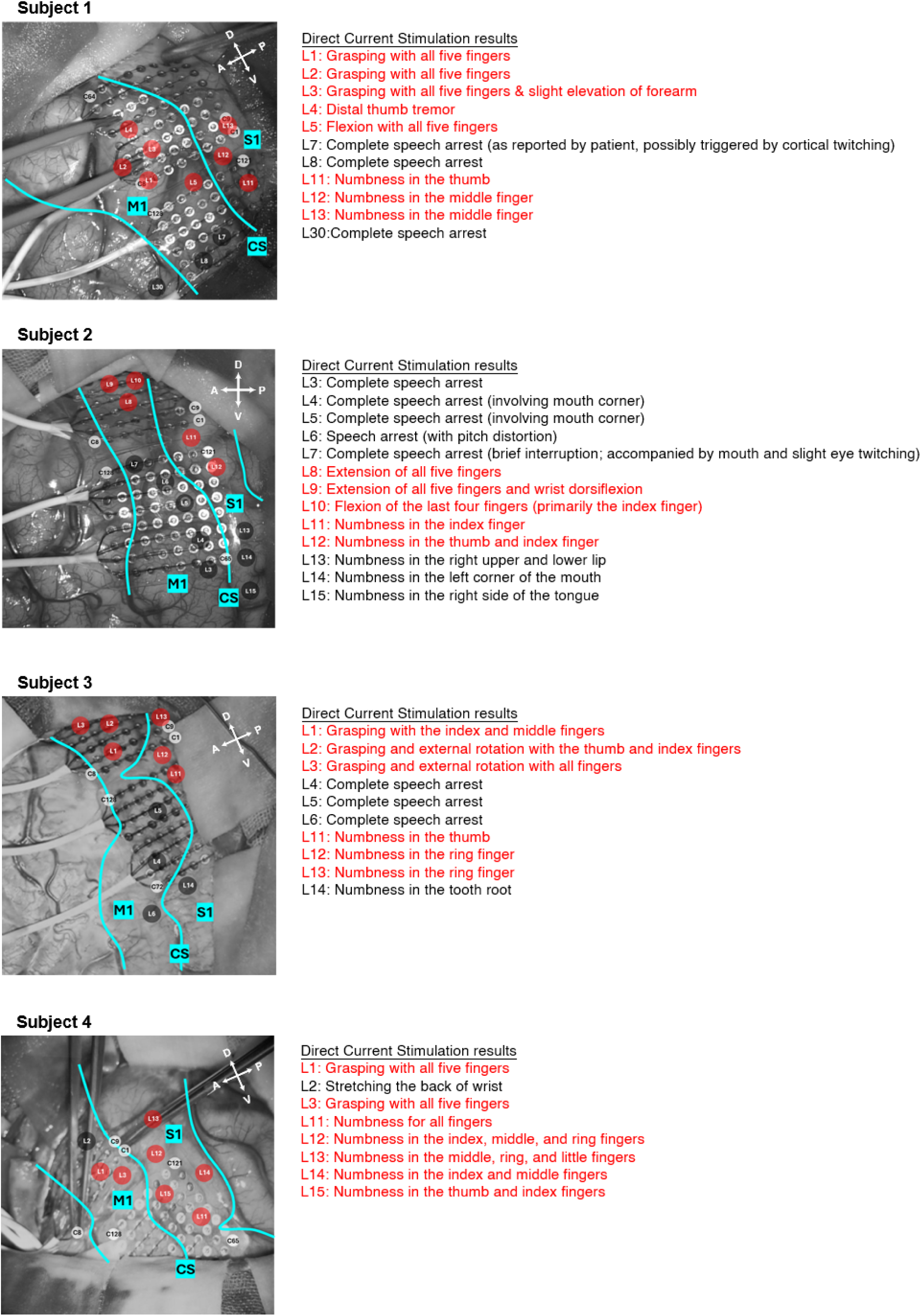
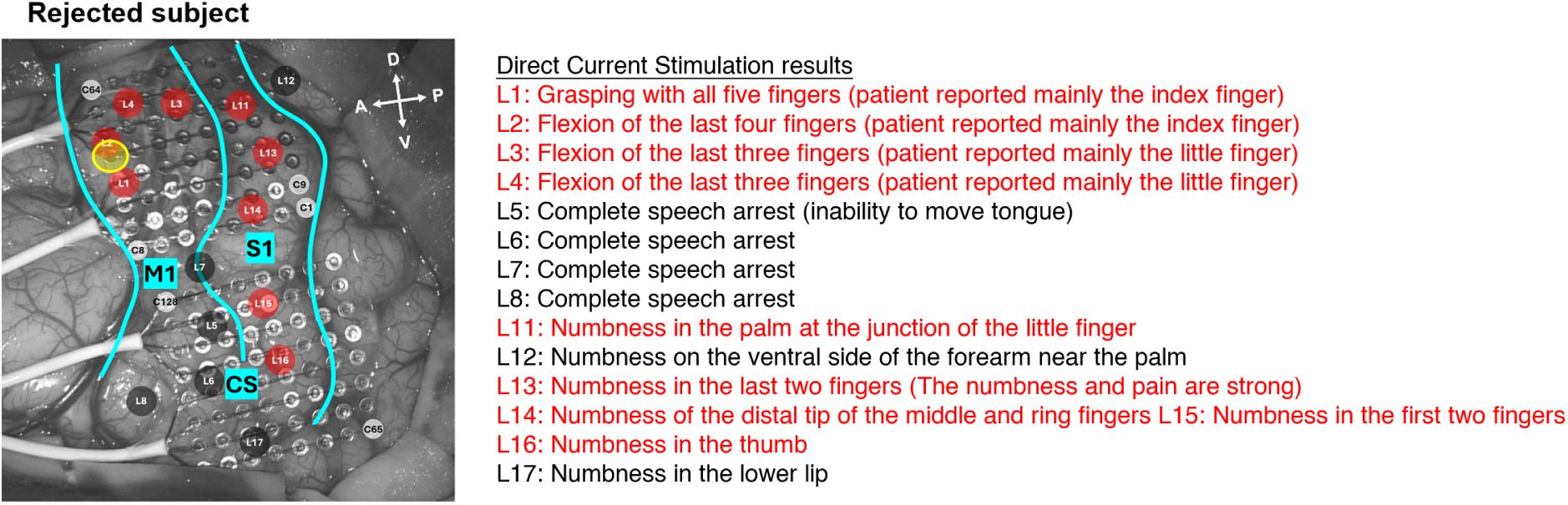
ECoG grids location. ECoG grids (8×8; two grids; 128 channels in total) placed over sensorimotor cortex in all subjects recorded. Sites where electrical stimulation evoked finger movement or sensation are marked in red; other responsive sites are shown in grey, with behavioral responses to electrical stimulation described on the right. The central sulcus (CS), primary motor cortex (M1), and primary sensory cortex (S1) are indicated. Orientation labels denote anterior (A), posterior (P), dorsal (D), and ventral (V).

**Fig. S2.**
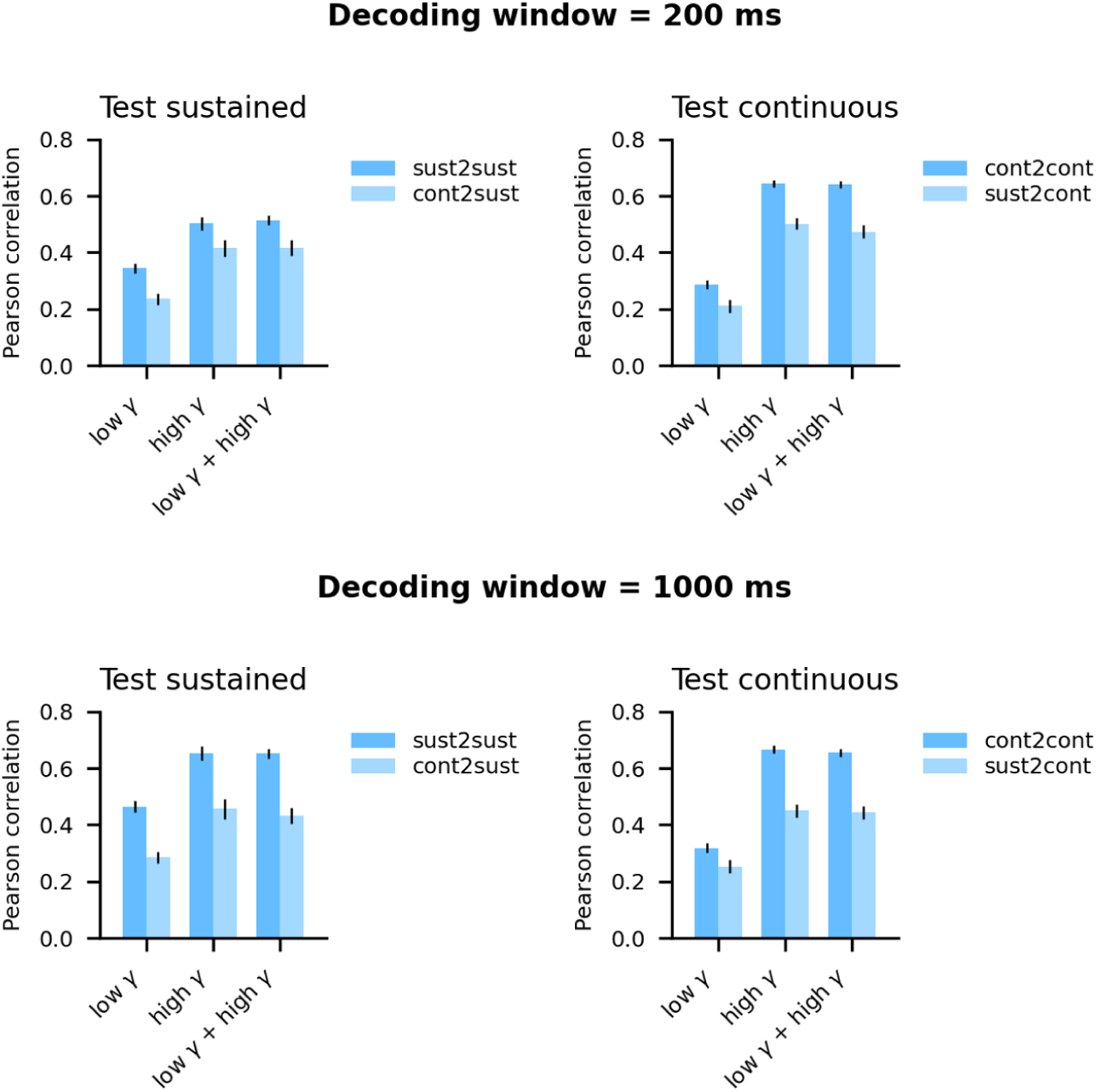
Contribution to decoding of low- and high-gamma. Pearson correlation between dataglove trajectories and the predictions of PLS in the seen and unseen conditions, trained with gamma features individually, and combined. Bars show the mean (± SEM) across all subjects and fingers (n = 20). High-gamma significantly outperforms low-gamma in all cases (Wilcoxon paired test, FDR-corrected, p < 0.005). No significant differences are found between high-gamma and low- and high-gamma combined.

**Fig. S3.**
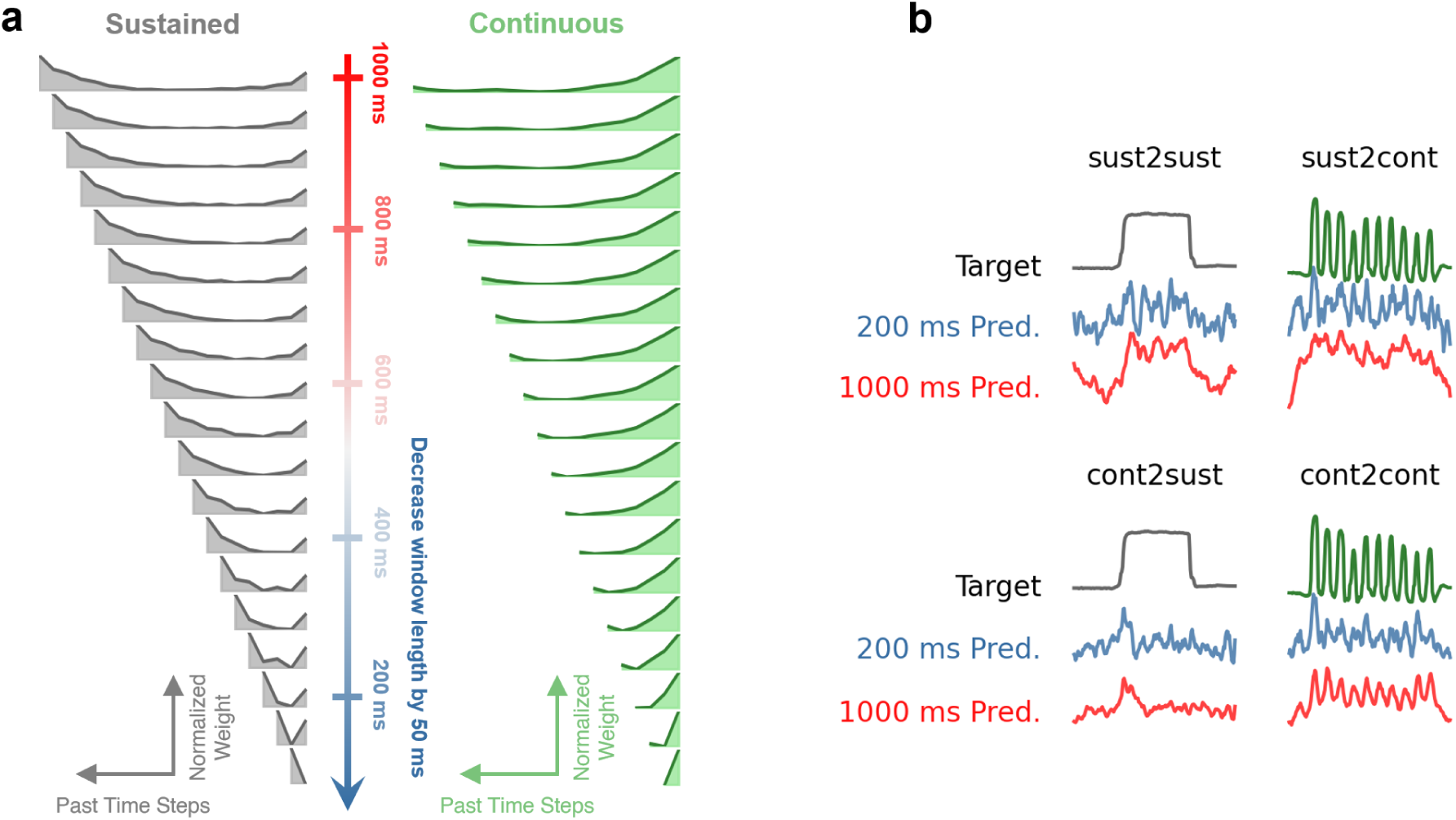
Short temporal windows enhance generalization. **a.** Decoder weights assigned to each time step within the window for PLS models. Left: models trained on the sustained task (grey). Right: models trained on the continuous task (green). Within each window, the leftmost taps represent earlier time points, and the rightmost taps represent recent time points. Larger amplitudes indicate greater contribution to the model’s prediction. **b.** Dataglove trajectories and corresponding PLS predictions of an example trial (subject 2, thumb) for short (200 ms, blue) and long (1000 ms, red) windows, shown for both the test sustained (left) and test continuous (right) task conditions.

**Fig. S4.**
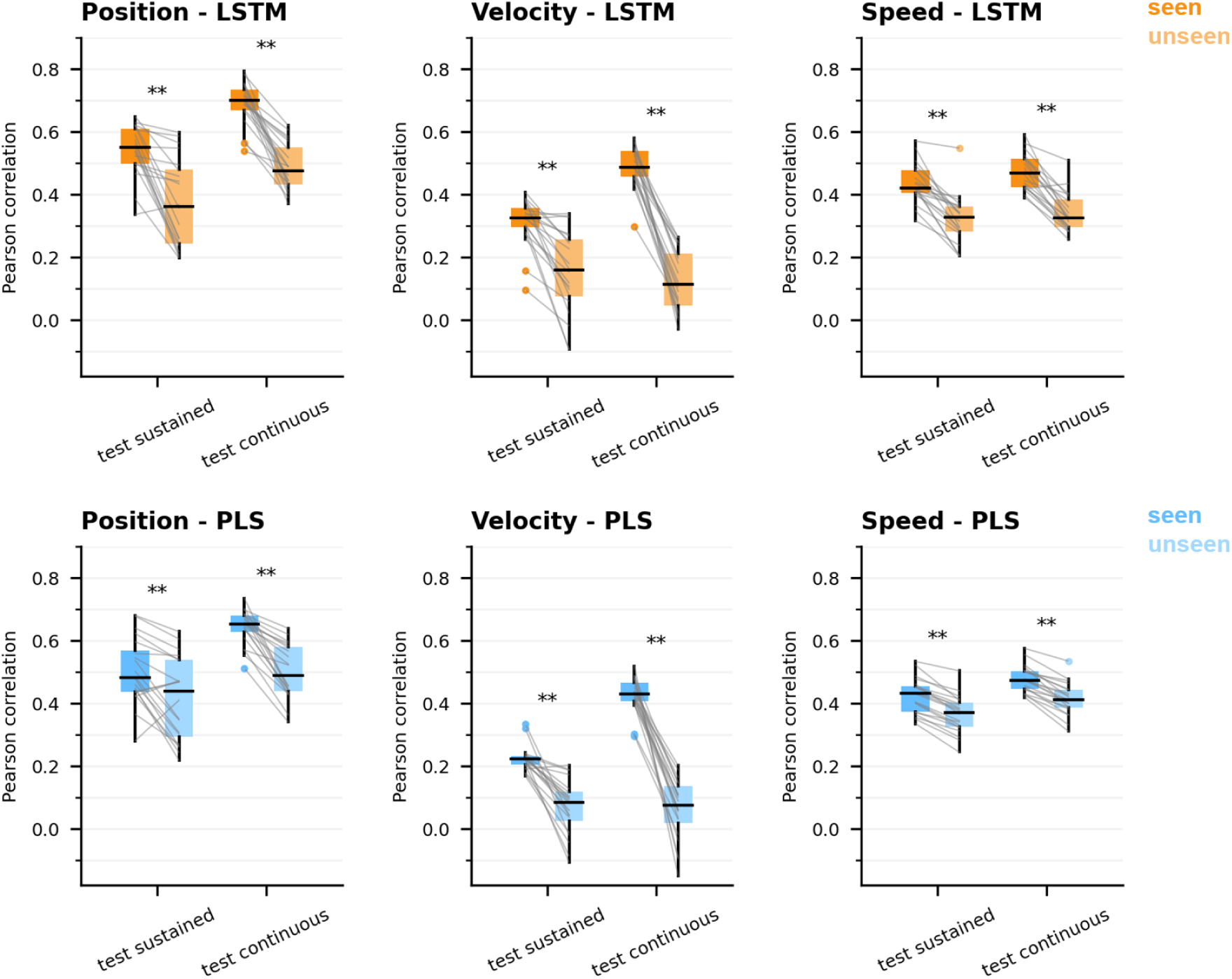
Comparison of position, velocity and speed generalization performance. Pearson correlation between target and predicted trajectories for LSTM (top) and PLS (bottom) in the seen and unseen conditions. Decoders are trained to predict finger position (left), finger velocity (middle), and finger speed (right, i.e. keeping only the magnitude of the velocity). High-gamma features and 200 ms windows are used in all cases. Boxes show the median and IQR range, whiskers show the most extreme non-outlier values (within 1.5 × IQR), and outliers are plotted as dots. Grey lines connect paired samples. Statistical significance is assessed with the Wilcoxon signed-rank test (**p < 0.005, FDR corrected).

**Fig. S5.**
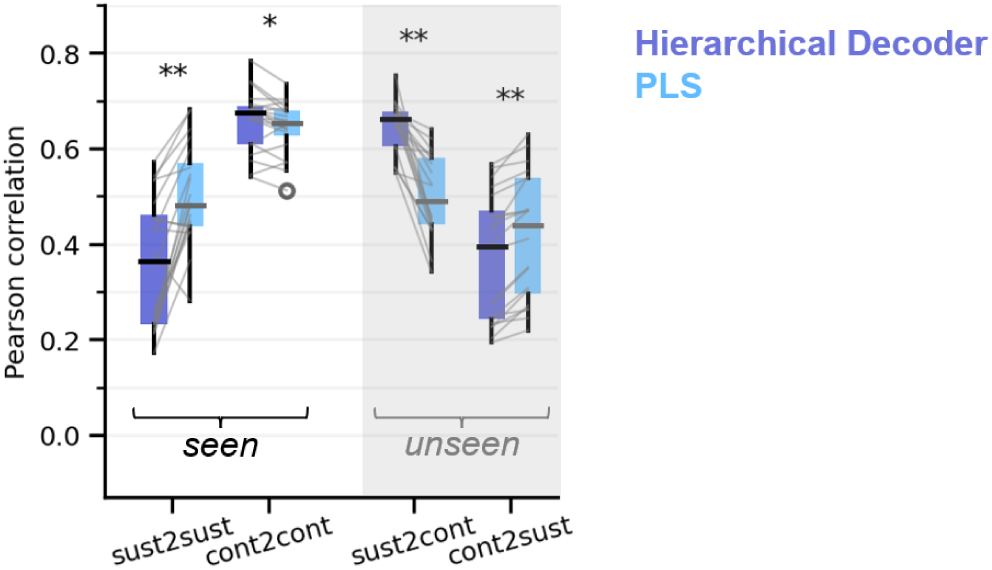
Comparison of a hierarchical decoder and PLS. Pearson correlation between target and predicted trajectories for a hierarchical decoder and PLS in the seen (left) and unseen (right, grey background) conditions. High-gamma features and 200 ms windows are used. The hierarchical decoder consists of two stages. First, a classifier distinguishes static from dynamic samples (logistic regression). Next, its output is used to weight the contributions of two PLS regressors, one trained exclusively on static data and the other on dynamic data. Boxes show the median and IQR range, whiskers show the most extreme non-outlier values (within 1.5 × IQR), and outliers are plotted as dots. Grey lines connect paired samples. Statistical significance is assessed with the Wilcoxon signed-rank test (**p < 0.005, FDR corrected).

**Fig. S6.**
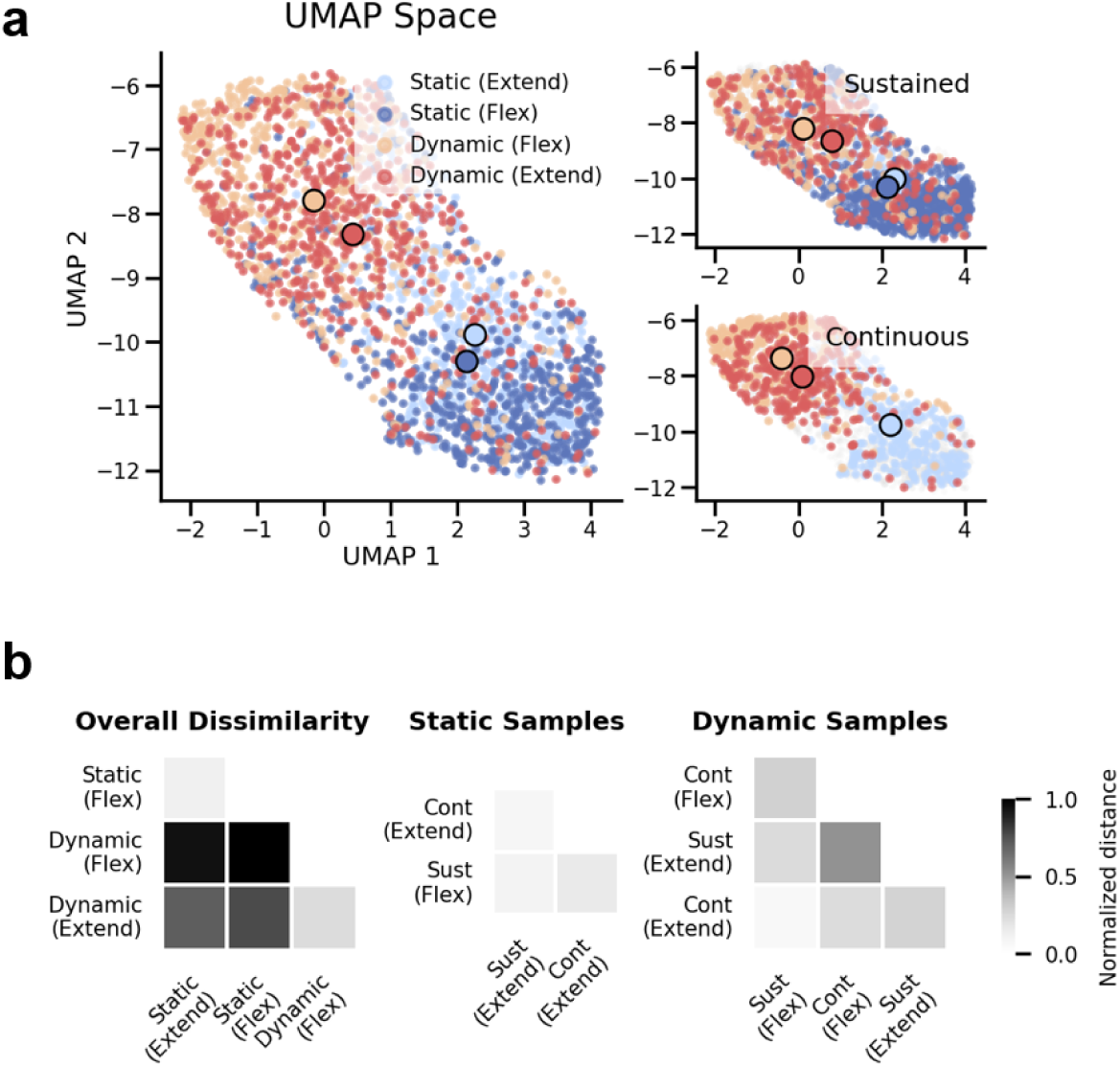
Neural differences between static and dynamic actions. **a.** Dimensionality-reduction of all samples considered in Fig. 5c using UMAP. Each point represents a high-gamma time sample, colored according to action type. Colored circles indicate the centroid of each action’s cluster. The right panels show the same PCA space when considering only samples from the sustained task or only samples from the continuous task. **b.** Dissimilarity between cluster centroids for the four action types, averaged across all subjects and fingers (n=20), for the UMAP space from panel a. The right panels show centroid dissimilarity when clusters from the two tasks are compared separately.

**Fig. S7.**
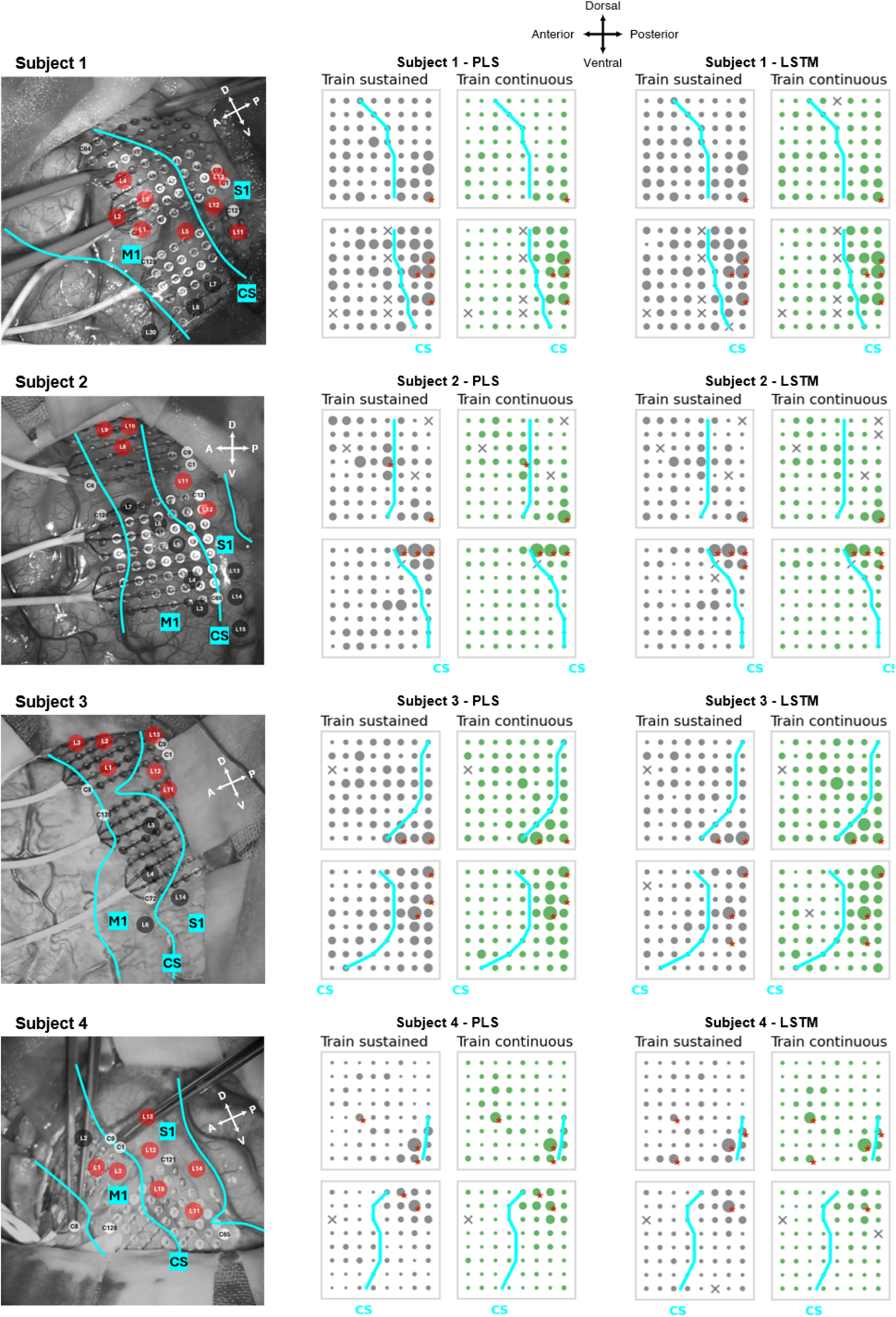
Channel wise contributions to decoders. ECoG grids anatomical location and channel-wise contributions to decoders (PLS: middle, LSTM: right) for all subjects. Larger circles indicate stronger contributions. Channels rejected during preprocessing are marked with an “x”. The central sulcus is shown in blue. The top five channels contributing most strongly across both tasks are marked with red stars.

**Fig. S8.**
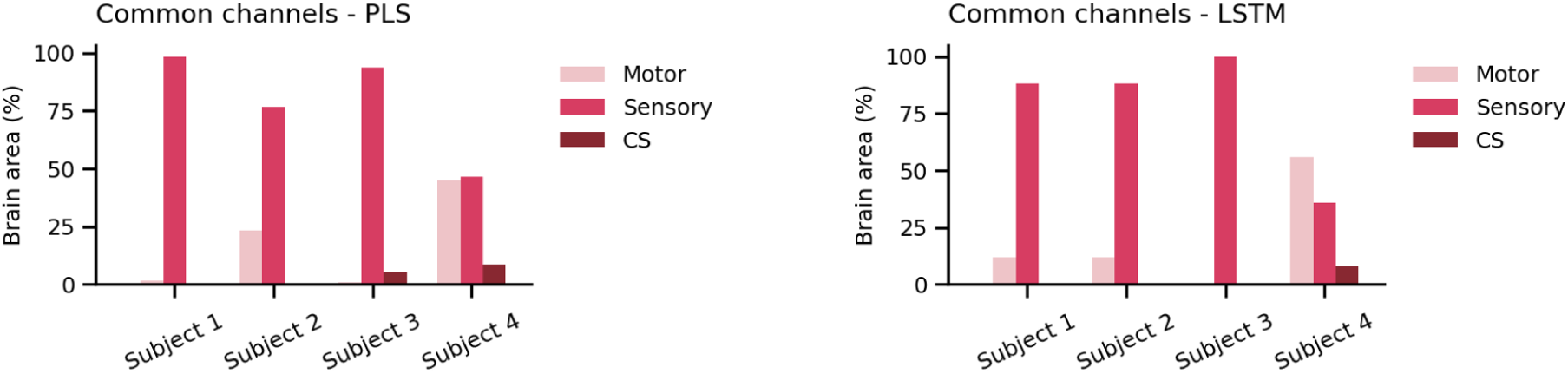
Anatomical distribution of common channels to the sustained and continuous tasks (motor: anterior to the central sulcus; sensory: posterior to the central sulcus; CS: channels directly over the sulcus) for PLS and LSTM decoders.

## Notes

### Competing Interest Statement

The authors have declared no competing interest.

